# Geometry of neural computation unifies working memory and planning

**DOI:** 10.1101/2021.02.01.429156

**Authors:** Daniel B. Ehrlich, John D. Murray

**Author notes:** Correspondence and requests for materials should be addressed to J.D.M.

## Abstract

Real-world tasks require coordination of working memory, decision making, and planning, yet these cognitive functions have disproportionately been studied as independent modular processes in the brain. Here we propose that contingency representations, defined as mappings for how future behaviors depend on upcoming events, can unify working memory and planning computations. We designed a task capable of disambiguating distinct types of representations. Our experiments revealed that human behavior is consistent with contingency representations, and not with traditional sensory models of working memory. In task-optimized recurrent neural networks we investigated possible circuit mechanisms for contingency representations and found that these representations can explain neurophysiological observations from prefrontal cortex during working memory tasks. Finally, we generated falsifiable predictions for neural data to identify contingency representations in neural data and to dissociate different models of working memory. Our findings characterize a neural representational strategy that can unify working memory, planning, and context-dependent decision making.

---

In time-varying environments, flexible cognition requires the ability to store and combine information across time to appropriately guide behavior. In commonly used delay task paradigms, a transient sensory stimulus provides information which the agent must maintain internally across a seconds-long mnemonic delay to guide a future response^1–3^. Working memory is a core cognitive function for the active maintenance and manipulation of task-relevant information for subsequent use. To guide flexible behavior, contents of working memory must interface with other cognitive functions such as planning and context-dependent decision making. Yet within neuroscience, these functions have been studied largely independently, and it remains poorly understood how the brain coordinates these processes in the service of goal-directed behavior.

Internal representations related to cognitive states can be revealed through recording neural activity during delay tasks in animals and humans. Neurons in the prefrontal cortex exhibit content-selective activity patterns during mnemonic delays of working memory tasks^4,5^. Selective delay activity during working memory is most commonly interpreted as representing features of sensory stimuli which can be processed to guide a later response. In the dominant conceptual framework, the proposed cognitive strategy thereby uses working memory representations that are fundamentally sensory in nature^4,5^. The sensory strategy for working memory has been challenged by observations of prefrontal delay activity that better correlate with diverse task variables, including actions, expected stimuli, and rules^2,3,6^. Furthermore, a growing literature characterizes substantial nonlinear mixed selectivity of features in prefrontal cortex, which can in principle support context-dependent behavior^7–9^. These diverse observations suggests a more unified framework is needed to account for the computational roles of working memory, planning, and context-dependent decision making.

Computational modeling has been fruitfully applied to examine potential neural circuit mechanisms supporting cognitive functions, including working memory and decision making^10^. Working memory functions are commonly modeled with distinct modular circuits, which do not account for mixed selectivity in prefrontal cortex and assume that working memory maintains sensory representations^10,11^. A complementary modeling approach utilizes artificial neural network models which are trained to performed cognitive tasks^12^. In contrast to hand-designed models, task-optimized recurrent neural network (RNN) models can perform delay tasks without requiring assumptions about structured circuit architectures or the form of working memory representations^13^. This approach is therefore well suited to examine computational mechanisms through which working memory representations can support flexible computations^14–17^.

In this study, we investigate the implications of a cognitive strategy in which internal states represent plans, rather than perceptions or actions. Specifically, our theoretical framework defines contingency states based on how future behaviors depend on upcoming events. We designed a task paradigm to dissociate contingency-based strategies from sensory- or action-based strategies for working memory and found that contingency strategy better explained human behavior in this task. RNN models trained on the task develop contingency-based solutions, and their internal representations capture diverse phenomena of neural activity during working memory. Lastly, we provide falsifiable predictions for neural activity to distinguish contingency representations from alternative computational schemes. Taken together, our study presents a theoretical framework for how working memory supports planning for temporally extended cognition and flexible behavior, which explains disparate behavioral and neural observations and is experimentally testable.

## Results

### Conditional delayed logic task

In many commonly used delay tasks, there are correlations among sensory stimuli, responses, computational demands or rules which prevent dissociable attribution of neural activity to specific cognitive task variables^1,18^. We therefore sought a task that involves (i) computation on informational inputs separated by a delay, (ii) exact intermediate computational states which re-occur frequently, (iii) trials for which the correct response can be predicted during the delay on some trials and not on others, and (iv) decorrelation of action, sensory, rule and computational state information.

To meet these demands we designed the conditional delayed logic (CDL) task, which applies a binary classification to two binary stimuli separated by a delay (**Fig. 1a**). We refer to the pre-delay and post-delay stimuli as “Cue A” and “Cue B”, respectively. The rules can be defined as Boolean logical operations, and one rule is presented tonically throughout each trial’s duration. On each trial, the agent is presented with a task rule, Cue A and Cue B and must generate an associated response. For example, in the OR rule the agent’s response should be “1” if either cue is equal to “1”. From 16 possible two-bit binary classifications, we will focus on 10 rules (**Fig. 1b**) for which the response is dependent on cues but independent of cue order. Rule, Cue A and Cue B are varied randomly across trials.

**Figure 1:**
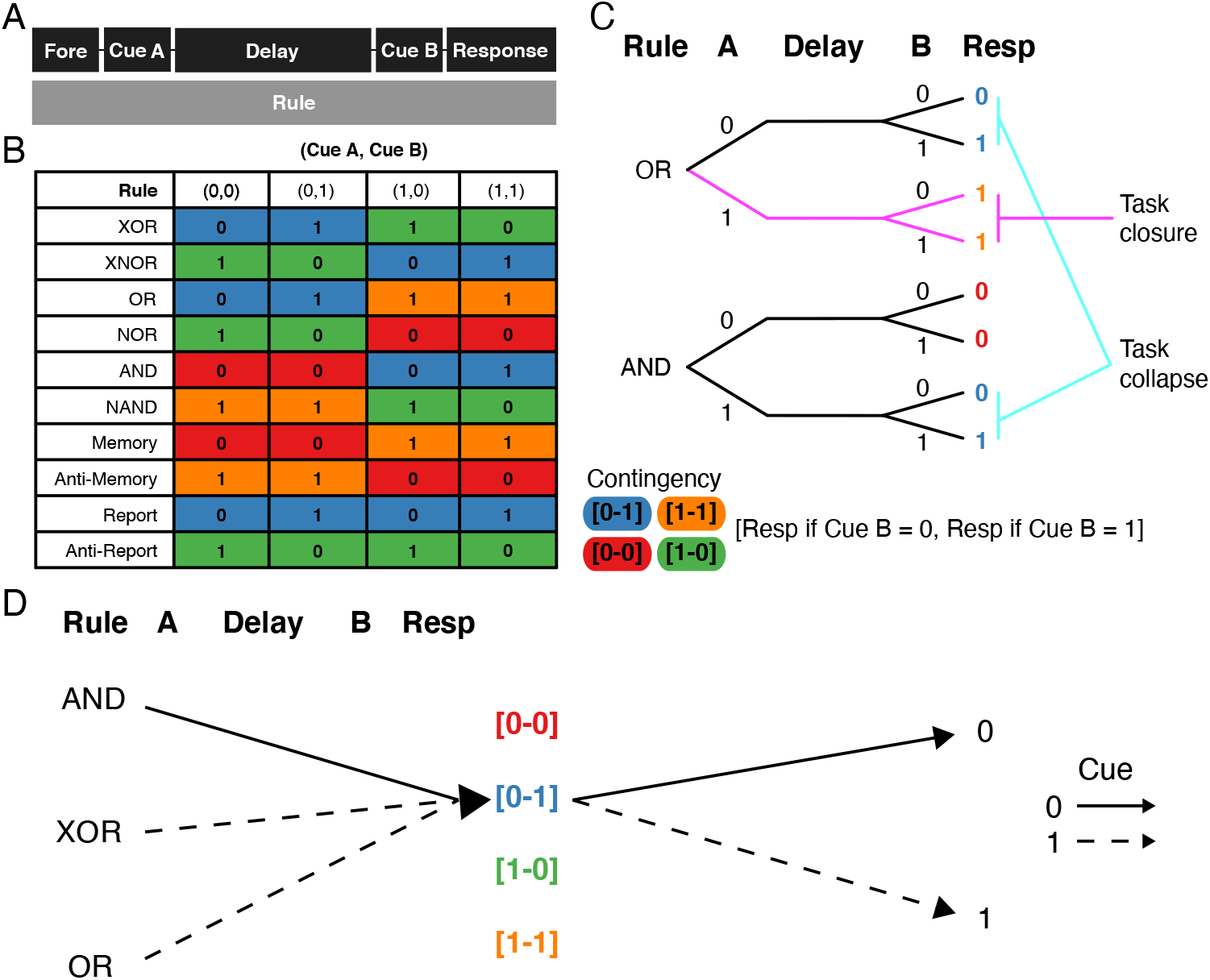
Conditional Delayed Logic (CDL) task and contingency states. **(a)** Time course of events in the CDL task. **(b)** The contingency table of responses by rule and (Cue A,Cue B) pair. Color indicates the contingency of that condition. **(c)** A schematic example of task closure and collapse between tasks and conditions. Top, tree structure of an OR trial, Bottom, tree structure of AND trial. The magenta lines indicates a condition (OR, Cue A=1) in which the trial is termed “closed” due to the fact that the correct response is independent of Cue B. The cyan lines indicate two conditions (OR, Cue A=0; AND, Cue A=1) which “collapse” during the delay. This is because after Cue A, their response patterns to Cue B becomes identical: in both cases if B=0 the correct response is 0, and if B=1 the correct response is 1. Color of responses indicate associated contingency. **(d)** Example solutions of the CDL task through contingency states. Solid lines represent a Cue of 0, and dashed lines represent a Cue of 1. All three tasks (AND, Cue A=0; XOR, Cue A=1; OR, Cue A=1) have a [0-1] contingency and therefore can be solved together after the delay.

### Contingency representations

To perform the CDL task, the agent must maintain a representation of task information across the delay. One might expect the agent to maintain the stimulus identity of Cue A, which would be sufficient to solve all trials. In this ‘sensory strategy’, which is commonly assumed in models of working memory, the task-relevant information is maintain across the delay to guide a response in conjunction with information in Cue B and rule^4,19^. An alternative ‘action strategy’ is possible for trial conditions in which the response can be preplanned during the delay^2,20^. Furthermore, the agent may represent the rule identity which is presented during the delay^3^. These representations are not exclusive.

In contrast to sensory, action or rule representations, an alternative strategy for the CDL task, and other working memory tasks, uses what we call ‘contingency representations’. Contingencies are defined by mappings from the upcoming cues to responses (**Fig. 1b**). For example, if the rule is OR and Cue A is “0”, then the contingency state during the delay is the mapping from Cue B being “0” or “1” to the correct response being “0” or “1”, respectively. Throughout this paper we will refer to contingency using the notation [R_0_-R_1_] where R_0_ and R_1_ are the response targets for Cue B being “0” and “1”, respectively. We would therefore say the above trial has a [0-1] contingency state, which could be represented in working memory.

The CDL task provides two important features that can be utilized by contingency representations: task ‘closure’, which gives insight into conditional computational demands; and task ‘collapse’, which enables identification of specific computationally relevant states. Closure describes the extent to which all information necessary to perform the next step of the task is already presented. In the CDL task, closure is defined as a condition in which the correct response can be decided from the rule and Cue A alone, and therefore the response can be pre-planned during the delay before Cue B. For example, if the rule is OR and Cue A is “1”, then regardless of Cue B the correct response will always be “1” (**Fig. 1c**). This enables the agent to plan the response directly following Cue A.

Collapse describes the condition in which the agent reaches the same computational branch point in a multi-stage problem independent of the prior path. In the CDL task, collapse describes when two different rule and Cue A conditions have the same contingency state with regard to Cue B. One example of this is in the OR and AND tasks: The [0-1] contingency state is reached both by the OR rule with Cue A of “0”, and by the AND rule with Cue A of “1” (**Fig 1c**). In this way two trials that share neither rule nor Cue A can share a contingency state, demonstrating functional collapse. As described later, this property will be crucial in dissociating contingency representations from sensory representations in neural delay activity.

### Computing through contingency representations

We found that one advantage of the contingency representation is that it reduces the overall classifier complexity problem of the task. The sensory representation requires 12 classifier hyperplanes to implement the ten rules, while the contingency representation requires only two (**Fig. 2a,b**). The contingency representation acts as a modular solution to the CDL task in which dynamics and representations are re-used in order to solve multiple conditions (**Fig. 2c**). Contingency representations solve the CDL task in two steps, a first step mapping from rule and Cue A to a contingency state and a second step mapping from contingency and Cue B to response. This decomposition turns a single complex classification problem into two simple ones. Interestingly, this complexity reduction can be formalized using a straightforward linear matrix decomposition between pre-delay features (rule and Cue A) and post delay features (Cue B and target) (**Supplementary Fig. 1**), which results in the contingency basis as described above (**Fig. 2b**).

**Figure 2:**
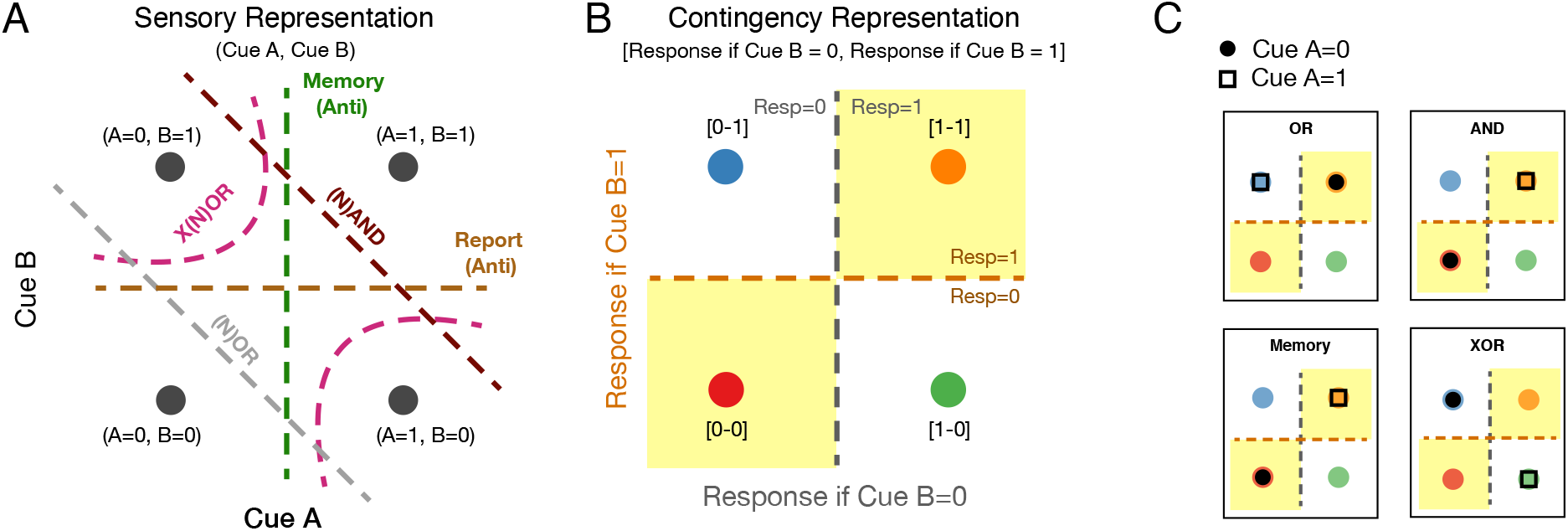
Geometric organization of contingency representations. **(a)** A schematic of the stimulus-based sensory representation of the CDL task. Each point represents a specific sensory combination (Cue A, Cue B) and each dashed line a separatrix that performs one of the rule operations in that space. Each line represents two rules as the negation of each rule is just the inverse mapping of the points on either side of the separatrix. **(b)** A schematic of the contingency representation. Each point represents a specific contingency state [response to B=0, response to B=1]. The dashed lines are separatrices capable of solving the ten tasks from the contingency representation. **(c)** An example of computing through the contingency representation for the OR, AND, Memory and XOR rules. Black circles(squares) represent the mapping of Cue A = 0(1) into the contingency representation for each task.

### Human behavior matches contingency strategy

While there are theoretical advantages to computing with contingency representations compared to sensory representations, it is unclear whether humans would use contingency as a strategy to solve the CDL task. To identify whether contingency is present in human behavior, we investigated the impact that closure and collapse had on response times (RTs) in a five-rule variant of the CDL task (**Fig. 3a**). We hypothesized that ‘closed’ trial conditions with task closure would elicit shorter RTs than ‘open’ conditions without closure, because on closed trials participants can pre-plan their motor response during the delay, whereas on open trials they must wait for Cue B to form their response.

**Figure 3:**
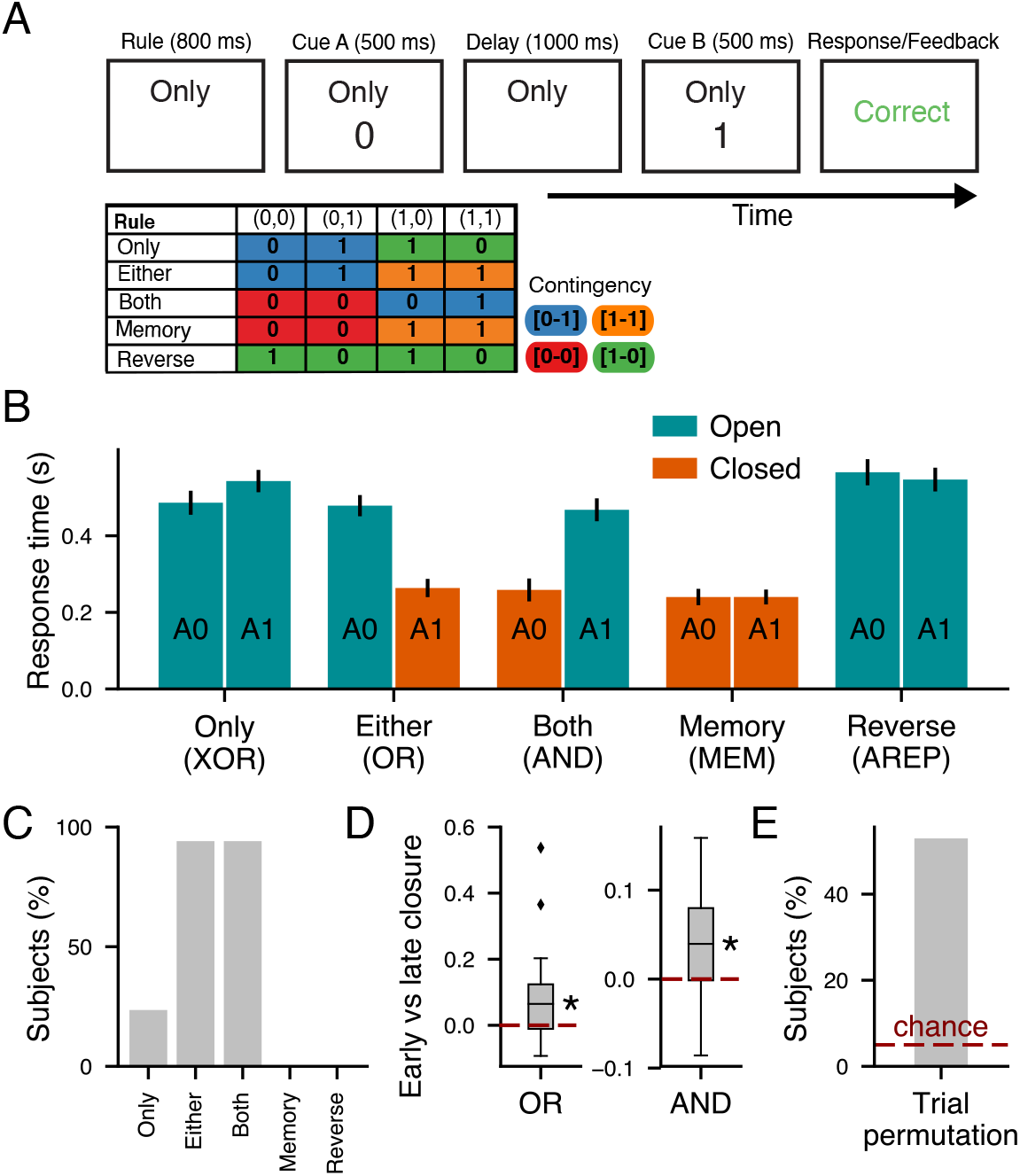
Human behavior on CDL task supports contingency-based strategies. **(A)** Top: The procession of stimuli shown to participants during an example trial (Only (XOR), (Cue A=0, Cue B=1)). Bottom: The table of responses by rule and (Cue A,Cue B) pair. Color indicates the contingency of that condition. **(B)** Mean response time by Cue A and rule. Error bars represent standard error of the mean across participants. Magenta bars are open trial conditions and orange bars are closed trial conditions. Logical rule names in parenthesis below labels. **(C)** Percent of participants with a significant (*p* < 0.05) difference in mean RT between Cue A=0 and Cue A=1 within a given rule. **(D)** The difference in closure effect, defined as the difference between the mean RT of the open condition and closed condition, between early trials (first two blocks of the task) and late trials (the last two blocks). Plotted separately for the OR and AND rules. **(E)** The percent of participants for whom contingency explained more RT variance than would be expected by chance for the linear model permutations. Dashed lines represent chance level, and stars indicate *p* < 0.05 (one-sided t-test).

We tested 17 human participants on the CDL task, measuring accuracies and RTs. Participants learned to perform the task with a high mean accuracy (97±1%) across all rules (**Fig 2**). To investigate cognitive and computational strategies underlying performance of the CDL task, we measured and compared RTs across task conditions both between rules and between Cue As (**Fig. 3b**). We found that RTs varied strongly across trial conditions and broadly sort into two groups according to closed vs. open conditions, with closed trials having a significantly shorter mean RT than open trials (*t* = −12.04, *p* = 10^−7^).

Crucially, the OR and AND subtasks, which vary on their closure status on a trial-by-trial basis, indicate that this RT difference between closed and open trials is not due to rule or cue differences. Within OR and AND, there remained a significant difference in RT between open and closed trials (OR; *t* = −6.4, *p* = 5 × 10^−5^, AND; *t* = −7.1, *p* = 2 × 10^−5^) (**Fig. 3b**). Within-individual analysis found that only for OR and AND was there a significant effect of Cue A on RT for the majority of participants (**Fig. 3c**), which would be expected if closure was driving RT. We observed that the closure effect, the impact of closure on RT, grew over the course of the session (OR: *p* = 0.018, AND: *p* = 0.009) (**Figs. 3d, 2d**), which suggests that the effect of closure can be learned through experience and be enhanced through training. The difference in RT between closed vs. open trials implies that they were capable of flexible updating of plans within a trial as they processed information from the rule and Cue A. This conditionally variant RT pattern is opposed to sensory strategies for which subtasks of similar complexity ought to take similar times to complete independent of how Cue A interacts with the rule (**Fig. 2a**).

While the analysis above indicates evidence against a sensory strategy for working memory, it does not specifically address whether participants use contingency representations with collapse across task conditions. To investigate the extent to which contingency is an explanatory task variable for human behavior, we used a linear modeling approach to measure the proportions of variance in RTs explainable by contingency, while controlling for effects of closure, response target, rule, and congruency between Cue B and response (see **Methods**). We utilized a permutation method to measure individual-level significance (**Fig. 3e**). We found that 53% of our participants (9 of 17) had significant explainable RT variance attributable to contingency, and that such a proportion was substantially higher than would be expected by chance (*p* = 3 × 10^−8^) (**Fig. 3f**). Collectively, our behavioral findings provide support that in the CDL task humans use contingency-based strategies and not sensory working memory strategies.

### Task-trained RNNs utilize contingency representations

To explore how contingency-based computations may be realized in a distributed recurrent circuit, we trained task-optimized recurrent neural network (RNN) models on the full CDL task (**Fig. 4a**). Our trained RNN model reached high accuracy across all subtasks (99.3% average accuracy, with all >95%). In order to identify the structure of contingency information in our trained RNNs, we used a simple subspace identification procedure^21^. Using the network state vector from the late-delay epoch (just before Cue B onset) we identified two contingency-coding axes maximally capturing variance in neural states across units in the RNN explained by the target response conditioned on each Cue B stimulus (see **Methods**). We then projected the late-delay state from each trial into this two-dimensional contingency subspace. This representation was found to cleanly and linearly separate between the four possible contingency conditions (**Fig. 4b**).

**Figure 4:**
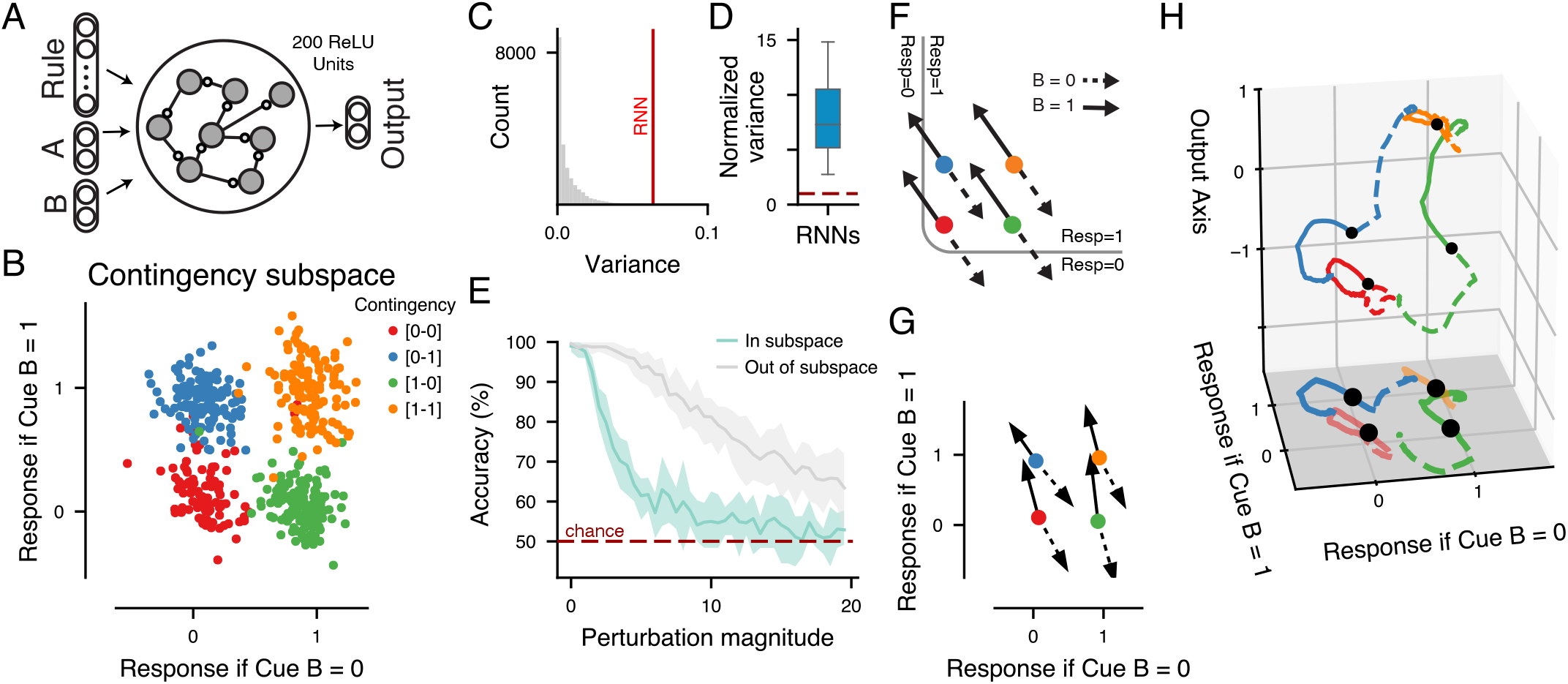
Functional contingency subspace in CDL-trained RNNs. **(A)** Schematic of RNN model structure. **(B)** We defined a contingency subspace through a linear regression of unit states during the late delay epoch to define two axes along which ‘Response to B=0’ and ‘Response to B=1’ were projected. The contingency subspace was identified on a train set of trials and we used held-out test trials for plots and analysis, for an example RNN. **(C)** A permutation test comparing the amount of variance in the contingency subspace compared to random alternative subspaces. **(D)** Analysis in B is replicated for 20 RNNs over random initializations and plotted is the distribution of variance captured by the contingency subspace compared to the mean variance of random subspaces of equal dimension. **(E)** Mean CDL performance (accuracy) across replicates for a perturbation analysis in which the RNN state was perturbed at the time point prior to Cue B onset by a random vector of a given norm either within the contingency subspace (cyan line) or orthogonal to the subspace (grey line). The perturbation magnitude, norm of the perturbation, was sequentially increased. Shaded regions represent the standard error of the mean across replicate RNNs. Red dashed line represents chance behavior. **(F)** Theoretical schematic of CDL response selection utilizing a contingency representation. Solid arrows represent trajectories caused by Cue B=0 and dashed arrows represent trajectories caused by Cue B=1. The grey separatrix boundary divides regions of state space in which the network will relax to a response of 0 and 1; **(G)** Mean contingency subspace trajectories of the example RNN model from onset to offset of Cue B. Trajectories divided by contingency and Cue B. **(H)** Three-dimensional mean RNN state-space trajectories from Cue B onset to trial end. X- and Y-axes represent the contingency subspace while the Z-axis is the output axis of the example RNN model (i.e., the difference between the magnitude of the two output units).

To test the importance of the contingency subspace, we measured the amount of trial-wise variance in late delay state vector captured by the contingency subspace and found greater than five times as much variance compared to random two-dimensional subspaces (**Fig. 4c,d**). We tested the functional relevance of the subspace for task performance through a perturbation approach. Just prior to Cue B onset, we shifted the state of the network by a given magnitude in a random direction which was either within the contingency subspace or orthogonal to the subspace. We found that RNNs suffered greater performance deficits from perturbations within the contingency plane, indicating the functional relevance of the subspace for the CDL task (**Fig. 4e**).

One advantage of the contingency representation is that it organizes delay representations such that they are already separable by mappings from Cue B to response. A single selection vector can therefore separate all states into the correct response (**Fig. 4f**). We analyzed the average displacement for trials in each of the contingencies during Cue B and found that they closely correspond to a theoretically optimal selection vector (**Fig. 4g**). The state-space dynamics following Cue B offset demonstrate a nonlinear consolidation trajectory from contingency to the appropriate response state (**Fig. 4h**), revealing that while the decision problem itself is linearized by contingency representations, task computation in the RNN relies on nonlinear dynamics.

Control analyses training artificial network models on CDL task variants investigated the computational properties leading to contingency representations. First, we trained an RNN on a task variant in which the rule is not presented until the Cue B epoch (**Supplementary Fig. 4**). This RNN perform this late-rule task with high accuracy, but it did not develop any contingency representations during the delay. This demonstrates that RNNs can solve the CDL task using a sensory strategy and that contingency representations are not an artifact of our analysis method. Second, we trained a three-layer feedforward network, in contrast to an RNN, providing task inputs into different layers of the network (**Supplementary Fig. 5**). Contingency coding only emerged as a dominant middle-layer representation when Cue A and rule inputs were provided upstream and Cue B downstream, which is the feedforward analogue of the temporal structure in the CDL task. These analyses suggest that computational stages of information processing, not specific choice of model architecture, robustly determine whether a contingency representation is formed by task-optimized network training.

### Model neural activity captures neurophysiological findings

While at the population level our trained RNNs can be structurally and functionally identified to be utilizing a contingency-based solution, it is unclear how that is instantiated at the level of individual units and therefore how it might relate to the findings in single-neuron recordings from animals performing working memory tasks. We profiled the selectivity properties of our unit activity vector (**Fig. 5a**). While the Cue A averaged traces showed substantial tuning within a rule, we found that between rules the apparent Cue A tuning would invert. For many units, however, this inversion across rules could be substantially explained by contingency. Using a linear model we identified the percent of variance explained in unit state by Cue A, rule and contingency. While we found that most units predominantly encoded Cue A during the stimulus epoch, by the late delay this pattern had changed to primarily encode contingency (**Fig. 5b**). Nonlinear dimensionality reduction via UMAP recapitulated the transition from sensory to contingency representations, and showed that contingency and sensory tuning are dominant factors driving unit states in the late vs. early delay epoch, respectively (**Supplementary Fig. 3**).

**Figure 5:**
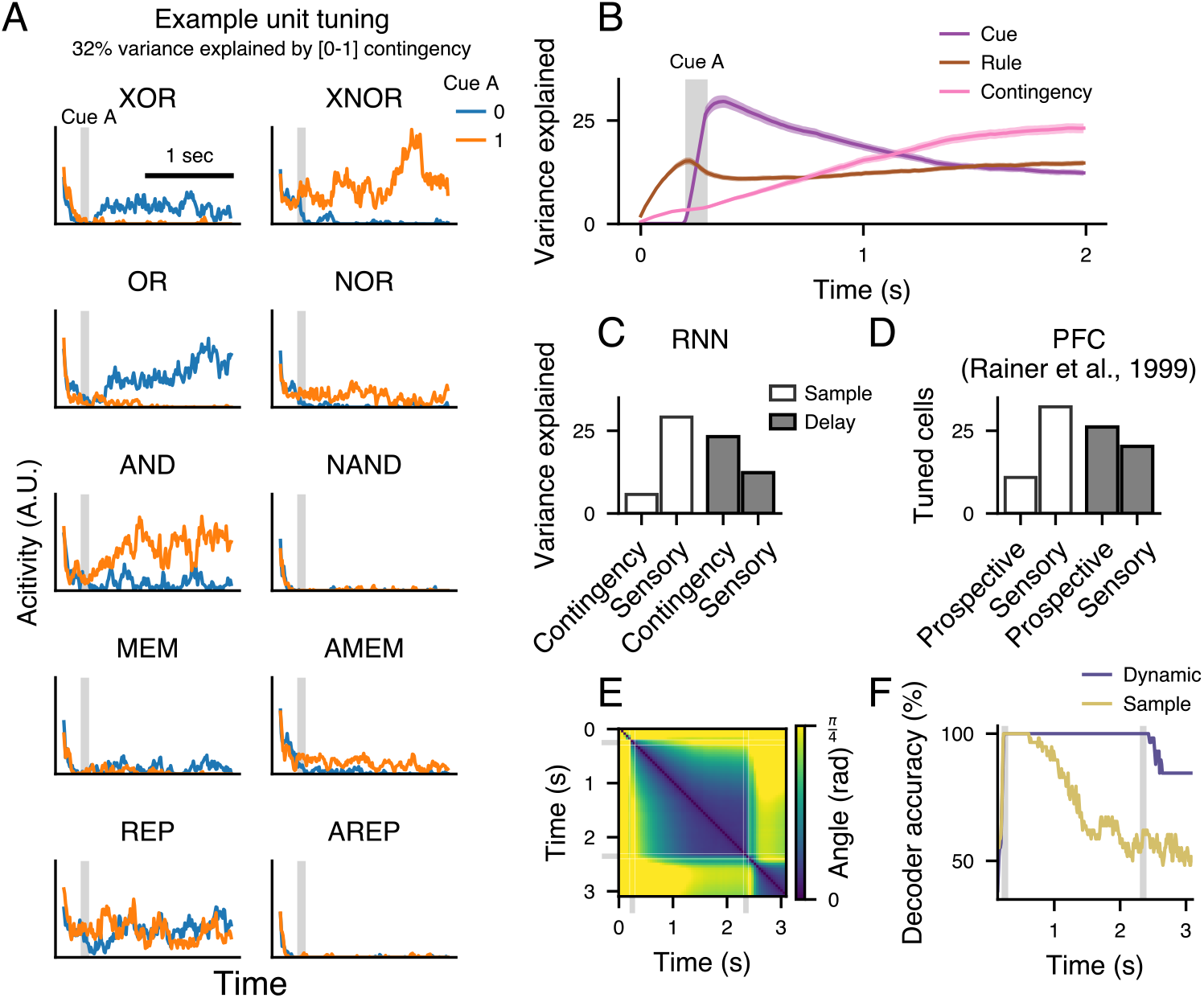
Contingency tuning captures neurophysiological responses in prefrontal cortex. **(a)** An example RNN unit with traces representing mean activity in trials divided by Cue A identity and rule. 32% of unit activity variance was explained by a binary predictor of whether it was a [0,1] contingency trial. Shaded grey region indicates the time of Cue A presentation. Traces plotted from trial onset through delay. **(b)** Mean percent of unit variance explained by rule, Cue A and contingency over time. Solid lines represent mean and shaded regions represent the standard error of the mean across replicate RNNs. **(c)** Fraction variance explained by prospective and sensory information for the sample (time point prior to Cue A offset) and delay (time point prior to Cue B onset) epochs in the example RNN model averaged across units. **(d)** Number of recorded neurons from monkey dorsolateral prefrontal cortex (PFC) significantly tuned to prospective and sensory information in the sample and delay epochs^6^. **(e)** Analysis of tuning dynamics across time. Activity in the example RNN was averaged within each condition and then PCA was used to identify the dominant axis of condition tuning. The angle between these axes were measured across each pair of time points. Gray shaded indicate Cue A and Cue B epochs. **(f)** A linear decoder was trained to read out Cue A identity from sample epoch RNN activity and tested throughout the delay epoch. This was compared to a “dynamic” decoder trained and tested on data from the same time point. Plotted is cross-validated accuracy for each decoder.

This transition in information coding during the delay matches prior analyses of single-neuron recordings from prefrontal cortex in monkeys performing working memory tasks. Rainer and colleagues analyzed neurons from alternating match-to-sample and paired-associate working memory tasks, and tested whether prefrontal neurons could better be understood as retrospectively tuned, wherein neuronal activity is tuned to sensory stimuli, or prospectively, wherein activity is tuned to expected future stimuli^6^. They found a transition from predominantly sensory coding during the cue epoch to predominantly prospective coding during the delay epoch. Our RNN units exhibit a matching transition from retrospective representation, tuned to Cue A, in the cue epoch to a fundamentally prospective representation, tuned to contingency, during the delay epoch (**Fig. 5c,d**).

This type of dynamic activity in persistent populations has drawn considerable interest in recent years, with many studies identifying a dynamic decoding axis of perceptual information from sample to delay epochs^22–24^. The contingency representation provides one possible explanation for this phenomenon. As during the delay the network shifts from a purely sensory to contingency representation if one were to only examine a single task it would appear that the axis along which Cue A is encoded has shifted. We used two analyses to detect dynamic working memory activity. In the first we evaluate change in the principal tuning axis over time using principal component analysis^24^ and in the second we train static and dynamic decoders showing a shift in the separatrix necessary to correctly classify the response from trial state vectors (**Fig. 5e,f**). In both cases our network matches phenomena previously interpreted as dynamic memory.

### Contingency subspace explains heterogenous neuronal tuning results

One question that arises from the above results is how contingency representations would be reflected in typical analyses which measure tuning of neural activity for task variables such as stimulus or rule identity. In the CDL task, tuning for Cue A or rule can be observed in contingency representations when the subtasks generate correlations between cue or rule information and contingency states. These correlations can exist even when Cue A, Cue B and responses are all independent. We examined this in CDL variants with only two rules. For example, the rule pair of Memory and XOR are such that contingency representations yield neural tuning to Cue A but not to rule, due to averaging over contingency states for those subtasks (**Fig. 6a**).

**Figure 6:**
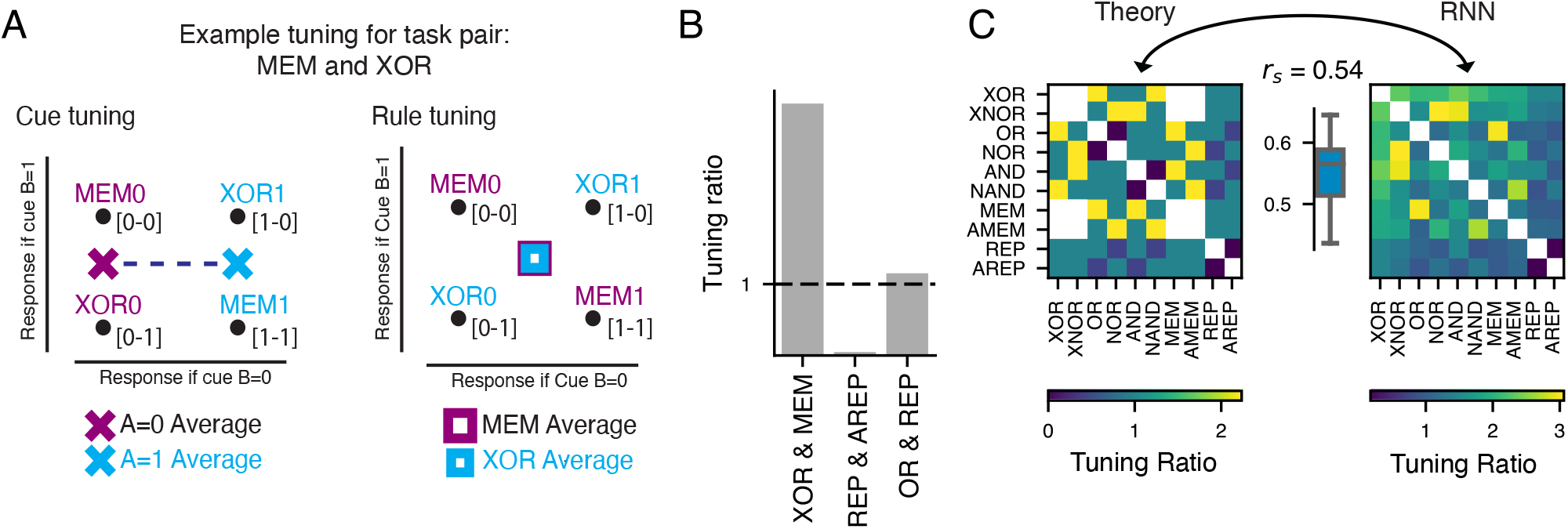
Contingency explains interactions of sensory and rule tuning for pairs of tasks. **(a)** Schematic of cue and rule tuning for an example pair of rules (MEM and XOR) utilizing a contingency representation. Task names are followed by a “0” or “1” indicating the Cue A on that trial and are placed in the appropriate contingency for that rule and Cue A pair. Left: X marks denote Cue A averages, with purple for Cue A=0 and blue for Cue A=1. Right: Square marks denote rule averages, with purple for MEM and blue for XOR. The dotted line in cue tuning indicates the difference between average cue states and therefor cue tuning. In contrast the overlap of rule averages indicates this task pair will show little or no rule tuning. **(b)** Cue-rule tuning ratio for three example pairs of rules. Bars show the measured tuning in the RNN when just that rule pair was analyzed. Dashed line represents equal cue and rule tuning. **(c)** Left: Contingency based predictions for cue-rule tuning ratio for each pair of rules in the CDL family defined as the Euclidean distance between cue averages and rule averages. A value greater than 1 indicates more cue tuning, A value less than 1 indicates more rule tuning and a value of 1 represents equal tuning. Right: Tuning ratio between Euclidean distances of cue and rule averages of state space representations of RNNs when analyzed for each pair of rules. Distances averaged across replicate RNNs. Inset: The Spearman correlation of (off-diagonal) predicted and measured tunings (*r_s_* = 0.54).

The nonlinear mapping from Cue A and rule to contingency can cause the same unit to be identified as rule- or cue-tuned depending on precisely which task the agent is performing (**Fig. 6a,b**). To demonstrate this we analyzed our network units separately for each possible rule pair in the CDL task, forming 100 possible pairs of rules. For each task pair, we could use the expected contingency states to determine theoretical prediction for cue and rule tuning (**Supplementary Fig. 6a,b**). We found that the theoretical contingency-predicted cue:rule tuning ratio significantly correlated with the ratio of tuning measured from the full CDL-trained RNN model in the late delay epoch (**Fig. 6c**).

Together these results show how linear analyses for neural tuning of contingency-coding units can yield apparent tuning to cues and rules. While in any real system, this simplified model of only contingency tuning will not fully explain all sources of variance, the extent to which even a simple model captures between condition tuning variance in our trained high-dimensional and nonlinear RNN demonstrates the robustness of the predictions made by contingency representations. Further, since tuning predictions can be made with small subsets of the CDL task it opens the door to experiments in animals for which it may not be feasible to train on a larger set of rules.

### Population coding reflects contingency in model network

To gain insight into how our theory of contingency representations differs from prior theories, and how they may be tested experimentally, we compared against two alternative models. The first is a pure sensory encoding model, in which states uniquely identify different sensory cues presented during the first stimulus epoch and the tonic rule input. The second is a randomly connected network (RCN) model, which generates high-dimensional linearly separable representations of rule and Cue A in order to do arbitrary classification problems^7,25^. These alternative model classes represent two ends of a continuum from most input structured representation (Inputs) to least structured (RCN). Further, they act as important controls because both models have been used to represent prefrontal computations in the literature^7,8,26^.

Due to our focus on between-condition representational structure, one useful analysis is representational similarity analysis (RSA)^27^. RSA enables the abstraction of high dimensional data into a set of comparisons between conditions called a representational similarity matrix (RSM). We use Cue A, rule and contingency as theoretical inter-condition templates, e.g. in the Cue A template all conditions with the same Cue A should be similar (**Fig. 7c**). The correlation between RSMs and theoretical templates measures the extent to which activity is structured by those features^27,28^.

**Figure 7:**
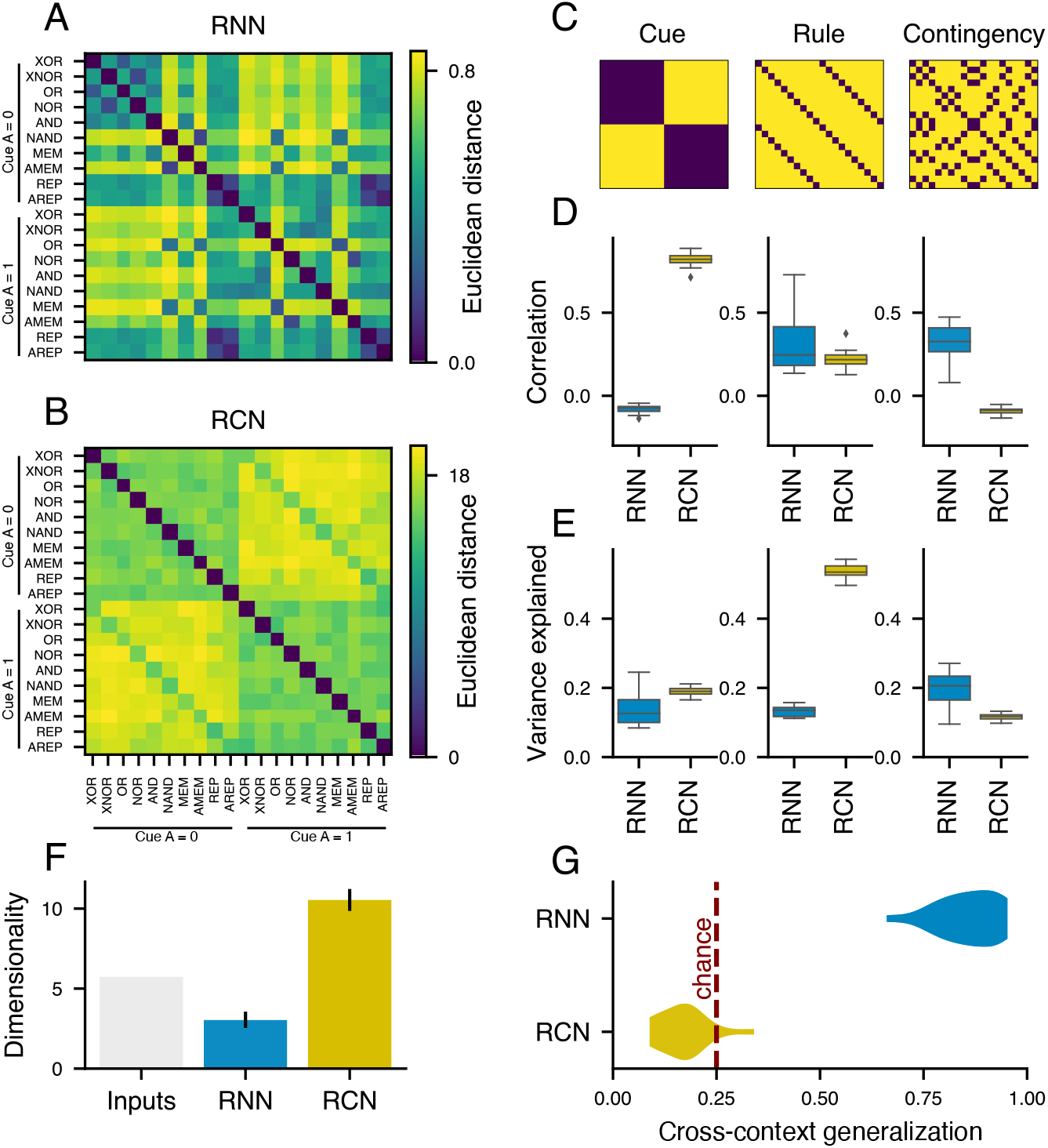
Model comparison and predictions. **(a, b)** Representational similarity matrices (RSMs) for **(a)** our example RNN model and for **(b)** a randomly connected network (RCN) model capable of performing the CDL task. Model similarity was calculated as the Euclidean distance between averaged late-delay epoch unit activity across conditions for the RNN model and mixed-layer activity for the RCN model. **(c)** Candidate RSMs for representational schemas organized by Cue A, rule and contingency. **(d)** Spearman correlation between the lower triangular portions of the candidate matrices and the RNN and RCN RSMs for each of our 20 replicate networks. **(e)** Mean fraction of variance explained by Cue A, rule and contingency across unit states during the late delay epoch, for the RNN and RCN. Plotted are the distribution of averages across replicate networks. **(f)** Dimensionality of late-delay activity of the RNN model, the RCN model and a “pure” model in which the network represents the Cue A and rule information orthogonally. Error bars represent s.e.m. across replicate networks. **(G)** Distribution of cross-context generalization (CCG) measured across replicate network for our RNN and RCN models. CCG was defined using a contingency subspace classifier. The contingency subspace (**Fig. 4b**) was fit as described above with one task condition held out. Then that condition was projected into the subspace and trials were classified based on the quadrant they fell in. Red dashed line represents chance classification.

The Inputs model by definition only contains input along the Cue A and rule dimensions, but we calculated RSMs for the task-trained RNN and the RCN model (**Fig 7a,b**). We then compared these observed similarity matrices to our candidate expected similarity structures constructed by Cue A, rule, and contingency (**Fig. 7c**). For the RCN model, Cue A predicts the most between-condition similarity, with rule explaining a lesser fraction and a small negative correlation with contingency. In contrast, the RNN model has inter-condition correlation best explained by contingency (**Fig. 7d**). These model predictions for population-level analyses were recapitulated in single-unit analyses by examining the amount of late-delay activity variance captured by a linear model including Cue A, rule, and contingency regressors (**Fig. 7e**).

To generate single-neuron predictions we fit a linear model for Cue A, rule and contingency to each unit’s activity in the late memory epoch to determine what fraction of variance in activity each regressor explains. The sensory model, by definition has single unit tuning towards Cue A and rule. The RCN model has a majority of units tuned to rule followed by Cue A with few units having substantial variance explained by contingency. The RNN model has the most variance explained by contingency followed by rule and Cue A (**Fig. 7e**).

The dimensionality of model state representations provides a complementary view into their geometric structure^29–31^. We found that the three models make starkly different predictions regarding the dimensionality of the delay representations. The sensory model has dimensionality governed by the inputs directly, and the RCN substantially expands that dimensionality. In contrast, the RNN contracts representations into a lower effective dimensionality which is only slightly higher than the theoretical minimal number of dimensions required by the task (**Fig. 7f**).

While the RCN model is designed for information to be generically decodable for a diversity of possible tasks, its high-dimensional representations are relatively unstructured and therefore will not generalize across conditions. We utilized a cross-conditional generalization (CCG) analysis^32^ to measure the extent to which the geometry across rule by Cue A pairs was preserved in both our RNN and RCN models. We first fit a contingency subspace, with all trials of a given task condition (rule × Cue A pair) held out. We then projected the held-out trials into the subspace, and defined a correct classification as a trial falling within the correct subspace quadrant. We found that our RNN models had a mean CCG of 86%, substantially higher than chance (*p* = 7 × 10^−19^, one-sided t-test), whereas our RCN models had a mean CCG score of 17% which does not exceed the chance level of 25% (**Fig. 7g**).

### Testing contingency coding in neural data

The analyses above are sufficient to characterize contingency encoding data generated for a known model class. We next examined how to statistically test for the presence of contingency coding in neural data in which we cannot know the process by which representations are generated, e.g. in experimental datasets of neural recordings. Contingency coding generated by a specific form of nonlinear interaction between task variables (e.g., rule and cue). To avoid distributional assumptions about a neural dataset, we devised a non-parametric partitioned variance approach to test whether a given dataset has more contingency-based nonlinear structure than would be expected based on the amount of other nonlinear structure in the data (**Fig. 8a**).

**Figure 8:**
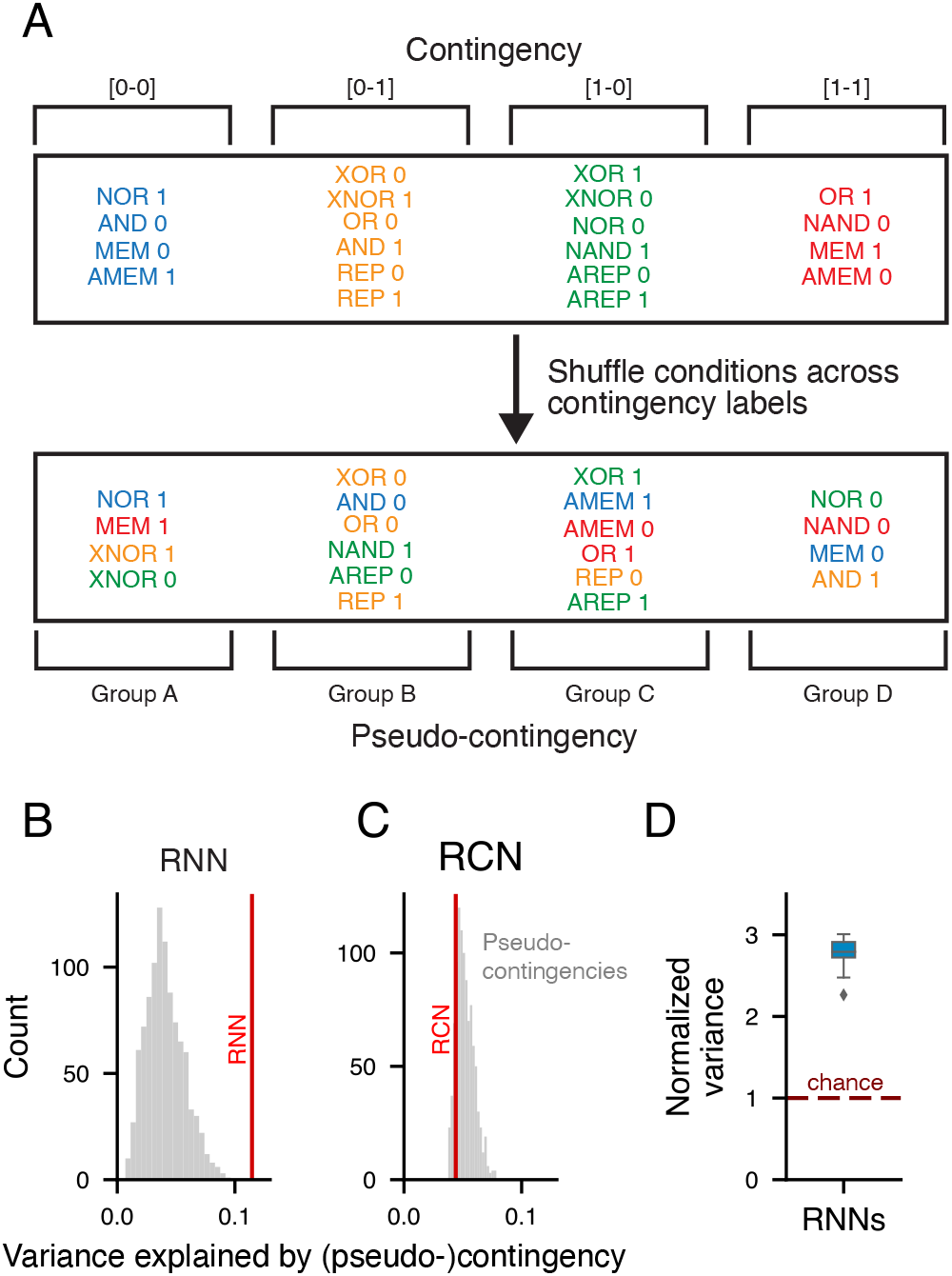
Partitioned variance analysis to test contingency coding. **(a)** Schematic of structure preserving contingency shuffling method. We construct pseudo contingencies of equal size to our actual contingencies but with random conditions. **(b,c)** Reshuffling test for proportion of contingency explained compared to shuffled pseudo-contingencies from **(a)**. The red line represents the mean variance explained by contingency for unit states in the RNN(RCN) model, while the grey bars represent the same measure for pseudo contingency replications. The RNN had significantly more variance explained by contingency than would be expected by chance while the RCN model did not. **(d)** Ratio of variance explained by contingency compared to mean pseudo-contingency shuffling, over 20 replicate RNNs. Dashed line represents chance level.

By shuffling contingency mappings across conditions (here, rule-cue pairs) into “psuedo-contingencies” and testing their ability to explain mean unit state variance, we can build a null hypothesis for how well we would expect a nonlinear interaction similar in structure, but different in specific conditions, to contingency would explain the data. Comparing random permutations to the measured quantity from the actual contingency mappings test whether the dataset is better explained by contingency than would be expected by chance independent of the general nonlinearity, feature mixing, or noise structure in the representation. We validated this approach with our RNN and RCN models, finding that our RNN model was better explained by contingency than chance (p<0.001) while our RCN model was not (**Fig. 8b,c**). Each replicate RNN model exhibited at least twice as much mean unit variance explained by contingency than the mean of its pseudo-contingency null distribution (**Fig. 8d**).

## Discussion

In this study we defined a representational schema for delay tasks in which network states encode the mapping between expected stimuli and actions termed the contingency representation. We developed a novel task, the conditional delayed logic (CDL) task, capable of dissociating contingency representations from stimulus or response representations. Human participants performing the CDL task demonstrated inter-conditional RT variance consistent with a contingency based strategy, and largely inconsistent with a stimulus memory strategy. In a trained recurrent neural network (RNN) we identified contingency representations and validated the functional role of contingency in the task. The structure of contingency representations in the model naturally captures experimentally observed phenomena including context-dependent tuning, mixed selectivity and dynamic coding for working memory. Lastly we generated falsifiable predictions for neural recordings of animals or humans performing the CDL task and compared results to alternative models.

Contingency representations can potentially unify memory, planning and cognitive control in a manner that does not require an external system to control the interactions between these subprocesses. In delay tasks where cues have a one-to-one correspondence with contingencies, the contingency representation is essentially indistinguishable from a sensory representation. In tasks where responses can be directly inferred from pre-delay cues, contingency states divide along future motor responses. Interestingly, tasks such as the CDL can exhibit both of the above cases. This directly links working memory to cognitive control through representational structure rather than positing an exogenous controller of inputs as in prior modeling^33^. Our behavioral data shows that humans can update their internal plans on the fly, between conditions of open contingency and closed contingency.

One intriguing property of contingency representations is that they can help to unify differing interpretations of neural activity in working memory delay tasks. Prior studies using neuronal recordings from primate association cortex have sought to dissociate retrospective sensory and more prospective signals for upcoming responses or expected stimuli^2,6,20^, and found tuning for task rules^2^. Using our RNN model, and top-down theory generated from the contingency subspace of the CDL task, we offer a putative explanation for some of the diversity of signals observed. By analyzing the network as it performed pairs of tasks with different geometric relationships in the contingency subspace, we found that units can appear tuned to rule, stimulus or action. This is due to the fact that the nonlinear transformation from rules and Cue A leads to different forms of collinearity in the contingency subspace.

The hypothesis that contingency could explain these results is bolstered by our behavioral results indicating that response times in our delay task are well explained by contingency. Response times have been shown to reflect cognitive processes^34^, including difference in preparation and intention^35^. RT measurements have been related closely to neural activity across cortex^36,37^. The observation that RT was strongly modulated by condition (closed vs. open) and that idiosyncratic RT variance could be explained by contingency in the CDL task provides evidence that participants utilized contingency during performance of the task.

Empirical analysis of unbiased samples of neurons have found that substantial populations are tuned to multiple features in a task. This phenomena has been termed mixed selectivity, with one hypothesis being that mixed selectivity generates a high dimensional and mostly unstructured basis to facilitate flexible computation^7,25^. Here we find that since many nonlinear combinations of rule and stimuli can lead to the same contingency state, units tuned to contingency can appear mixed. While we do find some units in our RNN are tuned to the Cue A and rule, units are consistently better tuned to contingency and therefore would appear as mixed to traditional linear analysis methods. This contingency tuning, however, is highly structured. Our finding is therefore better matched to recent research exploring the trade-off between generalization and flexibility as a function of between-condition structure^32^.

Recent studies have indicated there is substantial temporal dynamics in the pattern of activity seen during cue onset and early delay compared to late delay^23,38–40^. One hypothesis is that the mechanisms of neural persistence are intrinsically dynamic^41^. In our RNN, representations demonstrate what appears to be dynamics as the network moves from a stimulus dominant representation in the cue epoch to a contingency representation by the late delay epoch. In many experimental contexts contingencies are correlated with stimuli. In such cases the stimuli to contingency dynamics may appear as a change or rotation in stimulus coding (**Fig. 4**).

One important advantage to utilizing task-optimized RNN modeling in this study is that we do not impose any specific solution, thus allowing for emergent phenomena constrained by the task itself^13,16^. This strongly contrasts with most of prior working memory modeling in which the structure of delay states is pre-specified by the researchers^11,42^.

Further observability and controllability over the RNN during information processing enabled us to implement precise perturbations to measure the causal role of contingency representations in behavior. These features enabled us to expand on recent work identifying shared modules in a trained multi-task network performing many tasks previously identified with prefrontal activity^16^. In complement to the heuristic modularity they observed, the precision of the CDL task in sharing common computational intermediate stages allowed us to produce a top down theory of modularity that we could then use to guide investigation and measure exact modular overlap in our RNN.

Our study opens substantial theoretical and experimental questions for future research. One important question is whether differences in task demands would lead to alternative strategies to contingency being preferred. For tasks like CDL in which stimulus dimensionality is higher than response dimensionality, the overall complexity of the problem can be reduced by computing through contingencies rather than stimuli. For tasks with higher response dimensionality than stimulus dimensionality, however, it is unclear whether contingency representations would be advantageous. Furthermore, contingency is not always invertible, and therefore may be unsuitable in contexts in which it is often necessary to return to the previous cue. Of note, while contingency does reduce the CDL problem complexity, it does not reduce the amount of information stored mnemonically. To study these issues, the CDL task could be generalized to arbitrary dimensionalities of cue and action spaces, to systematically examine how capacity in both memory and problem complexity impacts strategies in human participants and in task-optimized artificial neural network models.

Future experiments can record neural activity patterns from human participants or animals performing CDL tasks, to test whether internal representations are governed by contingency^27,43–45^. Such an experiment could also help localize representational heterogeneity among cortical regions during the CDL task. Lastly, experiments could explore how changes in information flow between sensory, frontal and motor areas accompany the differential information routing for which we found behavioral evidence^46^. In turn, future modeling can incorporate aspects of known neurobiology which are potentially important for neural circuit computation, to extend beyond the relatively simple architecture used here. Neurobiologically motivated properties such as Dale’s principle^47,48^, short-term synaptic plasticity^49^, multi-regional connectivity constraints^50^, or attentional mechanisms could be explored for their effects on the emergence or structure of contingency representations.

In conclusion, we introduced a representational schema, contingency representations, capable of unifying working memory and planning without use of an external controller. We found evidence of these representations in human behavior. In a task-optimized neural network model we identified ways in which this representation can explain results on tuning from neurophysiological experiments on working memory, as well as provide new testable hypotheses for future studies of neural activity during cognitive tasks.

## Acknowledgements

We thank members of the Murray Lab for useful discussions and Rishidev Chaudhuri for feedback on an earlier version of the manuscript. This research was supported by NIH grant R01MH112746 (J.D.M.) and the Gruber Foundation (D.B.E.).

## Author Contributions

D.B.E. and J.D.M. designed the research. D.B.E. performed the research. J.D.M. supervised the project. D.B.E. and J.D.M. wrote the manuscript.

## Methods

### Behavioral task description

On each trial of the behavioral CDL task, the participant is shown two transient binary stimuli separated by a delay in addition to a rule cue on throughout the entirety of the trial. The participant is tasked with using the rule and stimuli to determine and report the correct response (**Figs. 3a**).

To increase statistical power and reduce training difficulty, five rules were selected from the ten original CDL rules (**Fig. 3a**): OR, AND, XOR, Memory (MEM) and Anti-Report (AREP). The OR, AND and XOR subtasks follow the stated Boolean logical operations. For the MEM subtask, the participant reports the identity of the first stimulus, Cue A (i.e., the stimulus that appears before the delay). For the AREP subtask, participants report the opposite of the second stimuli, Cue B (i.e., the stimulus that appears after the delay). For ease of understanding, and to reduce effects of differences in familiarity with logical operations, we displayed the five rules to participants as “Either” (OR), “Both” (AND), “Only” (XOR), “Memory” (MEM), and “Reverse” (AREP) rather than their logical labels (**Fig. 3a**).

The five rules selected for the behavioral task were chosen such that there were 2 conditions each of [0-0] and [1-1] contingency, and 3 each of [0-1] and [1-0] contingency. This helped ensure no imbalance in the proportions of trials between the two closed contingencies nor between the two open contingencies. Further, since there was one more condition for each of the open contingencies the shorter response time for open trials could not be explained by frequency.

Preceding Cue A was a foreperiod of 800 ms in which the participant was shown only the rule cue in the upper center of the screen. The rule cue would then remain on the screen throughout the entire trial until feedback. Following the foreperiod, the participant was shown Cue A, as either a “0” or “1”, in the center of the screen below the rule cue, for 500 ms. After Cue A was removed, only the rule was visible during a delay of 2000 ms. Cue B was then shown, as either a “0” or “1”, in the same location as used to display Cue A. The participant was able to respond by key stroke, with either a “left arrow” for “0” or a “right arrow” for “1”, at any point following Cue B onset. If the participant failed to make a choice within 2000 ms after Cue B onset, the trial would end and the participant would see a message telling them they timed-out, for a duration of 3000 ms. If the participant responded they would see either a “Error” or “Correct” feedback on the screen for 500 ms before the next trial begins. For each trial, rule, Cue A, and Cue B were selected randomly and independently.

The response time (RT) was calculated as the time between Cue B onset and key stroke response. Prior to the task, participants were instructed to respond as “quickly and accurately” as possible, and that they could respond as soon as the second stimulus (Cue B) appeared on screen. In order to control for any bias either in response speed for “left arrow” vs. “right arrow” or bias in response to “0” and “1” stimuli, participants were randomly assigned a binary “key mapping variable” which would either run the task as described above, or swap response mappings (i.e., left vs. right arrows to report responses) as well as “0” and “1” stimuli mappings.

Each participant completed 6 blocks of 64 trials each. The task was implemented through the PsychoPy package^51^ in Python, and run on a Macbook Pro laptop.

### Behavioral collection

18 participants were recruited from the general population between the ages of 18 and 35 in the New Haven, CT area. All participants completed and signed an informed consent approved by the Yale Institutional Review Board (IRB) and were paid for their participation. We used flyers and online advertising as our main recruitment methods. Of our 18 participants, 11 were female. The mean age was 24.3 years (standard deviation 4.2 years). Participants on average had 17.4 years of education (standard deviation 2.6 years).

Participants who contacted the lab were given a brief phone screening for eligibility. Participants with psychiatric diagnosis or history were excluded from the sample, as well as participants who had substantially impaired and uncorrected eyesight and participants with less than a 5th-grade reading level.

Eligible participants were invited into laboratory testing room, asked to read and sign an experimental consent form and then a brief demographic survey prior to task training. Following this they were shown a brief task description that explained the task, including the order of events, the goal, and the task. Participants were informed that their reward would be a function of accuracy and response time but were not told an exact calculation for reward.

Participants were then given a training block of 40 randomly selected trials of the CDL task. Participants who failed to get greater than 90% accuracy were required to repeat the training block. If the training block was failed three times the participant was paid the base compensation and excluded from the study. Only one participant was excluded this way.

Participants then performed 6 blocks of 64 trials, after each block the participant was able to take a self timed break. At the end of the task, their bonus reward was calculated based on their RTs and accuracy according to:

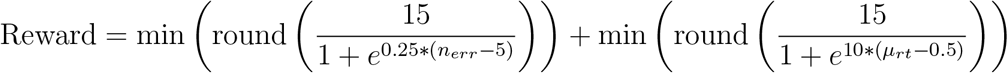

where *n_err_* represents the total number of errors and *μ_rt_* represents the average RT (in seconds) for correct trials. Reward was paid in US dollars at the end of the experiment, following a brief demographic survey.

### Behavioral analysis

To test the differences between closed and open trials, we averaged each participant’s RTs by closure, averaging all open trials and all closed trials, and then conducted a paired two-sided *t*-test across participants. Similarly, to test the significance of closure on conditionally closed trials, we averaged the RTs of each condition (Cue A × rule combination) separately. Then we ran a two-sided paired *t*-test between mean Cue A=0 RTs and Cue A=1 RTs across participants within each task. We tested the individual-participant significance by using a two-sample *t*-test to directly test the significance of the difference between Cue A=0 and Cue A=1 trials, for each rule independently, for each participant (**Fig. 3c**).

To examine learning effects, we averaged a given participant’s RT by condition (Cue A × rule) for the first two blocks which we termed “early” trials, and the last two blocks which we termed “late” trials. We determined the closure effect for OR and AND trials as the difference, for that participant, in mean RT in the open condition and the closed condition of those tasks. We termed this difference the closure effect. Then we calculated a two-sided paired *t*-test to measure whether this closure effect had increased from early to late trials, across the group (**Fig. 3d**).

To examine the effect of contingency on behavior, we measured the variance in trial-wise RT that is explainable by contingency. For each participant, we fit a linear model on correct trials, with RT as the predicted variable and the predictors being contingency, closure, rule, target, and congruency. Contingency was the contingency that would be applicable during that trial, based on the rule and Cue A. Rule is either XOR, OR, AND, Memory or Reverse (Anti-Report) depending on trial. Target was the correct response for that trial. Lastly, congruency is a binary variable representing whether Cue B was the same as the target. Congruency was used to control for possible effects of pro-/anti-match bias in RTs. Then we fit a second linear model with all the predictors above except contingency. Each predictor was encoded as a one-hot and models were fit with an L2 regularization penalty of 0.01.

Using these models we then calculated the percent of RT variance explained by each model. The difference between the full model and the model lacking a contingency predictor we termed the “ΔEV”, representing the amount of additional variance explained by accounting for contingency.

We tested for statistical significance using a permutation analysis, in which we shuffled trial contingency labels 1000 times and recalculated ΔEV, forming a null distribution against which we conducted hypothesis testing. 53% of participants (9 of 17) had a ΔEV greater than 95% of permuted samples. We then used a binomial test to identify the significance of the proportion of participants for whom contingency was found to be a significant predictor (**Fig. 3e**).

We calculated the variance attributable to contingency using the models below. Model 1 includes contingency as a predictor, and is specified by:

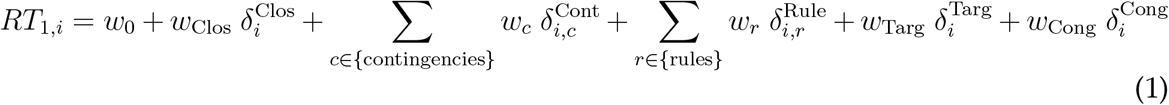

where *RT_i_* is the response time on trial *i*, and regression dummy variables are defined as Kronecker delta functions for the match of conditions in trial *i*. 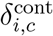 is 1 when the contingency of trial *i* matches contingency *c* and 0 otherwise. 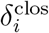 is 1 or 0 when the trial’s closure condition is closed or open, respectively. 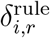 is 1 when the rule of trial *i* matches rule *r*. 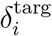 is 1 or 0 when the trial’s target condition is 1 or 0, respectively. 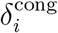 is 1 or 0 when the trial’s congruency condition is congruent or incongruent, respectively. Model 2 includes all predictors from Model 1 except for contingency:

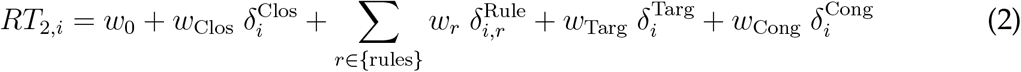

Here Model 1 has 12 parameters (1 constant, 4 for contingency, 1 for closure, 5 for rule, 1 for target, and 1 for congruency), and Model 2 has 8 parameters.

We quantified the sum of squared errors (SSE) for each model:

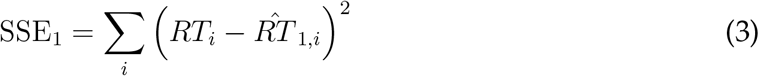

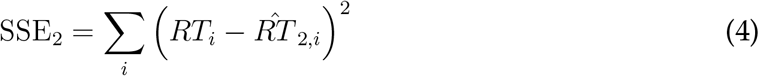

where the sum is over trials, and 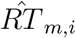 is the predicted RT for trial *i* from model *m*. The proportion of explained variance (EV) is given by:

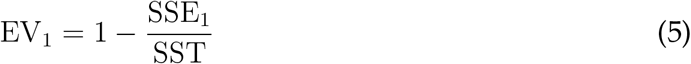

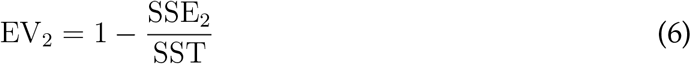

where SST is the total sum of squares (variance) of RT across trials. Then the amount of extra explained variance from including contingency regressors is ΔEV = EV_1_ − EV_2_.

### RNN model architecture

All RNN experiments were performed on continuous-time recurrent neural networks discretized to a 10-ms time step. RNNs contained 200 recurrently connected rectified-linear units (ReLU) in the hidden layer. Recurrent units received input as a linear combination of network inputs. Outputs were linear readouts of rectified recurrent unit activity. The network was governed by the following equations:

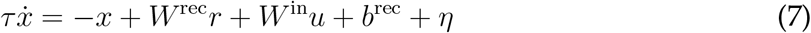

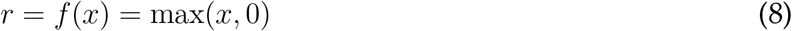

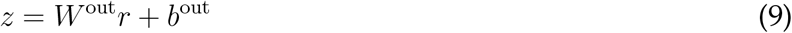

where *W*_rec_ is the recurrent weight matrix, *W*_in_ is the input weight matrix, and *W*_out_ is the output weight matrix. *x* is the unit state vector, *r* is the unit activity vector, *u* is the vector of inputs at a given time, and *z* is the output vector. *b*_rec_ and *b*_out_ are constant biases into the recurrent and output nodes respectively. *τ* = 100 ms is the internal time constant of units and *η* is gaussian noise added to the recurrent layer. Lastly, *f* (*x*) is the ReLU positive linear rectification function.

All inputs and outputs were represented using one-hot encodings. The network received cue input from 4 input channels, with the first two being assigned to represent a binary input during the first stimulus epoch, Cue A, and the latter two representing binary input in the second stimulus epoch, Cue B. The network received rule input through 10 channels with each channel associated with the instructed subtask being on and all others off^16,21^. The network response was recorded from two output nodes, each representing one of the two responses. Final response was determined by the output node with the greatest activity at the final time-point.

### RNN task description

On each trial the network received as inputs two binary cues and was tasked with choosing the correct response, as designated by a context cue presented tonically through the task. The X(N)OR, (N)OR, and (N)AND tasks followed the logic of their rules as applied to binary stimuli. For the Memory (MEM) and Anti-Memory (AMEM) tasks the network had to respond with the first cue or the opposite of the first cue, respectively. Similarly, for the Report (REP) and Anti-Report (AREP) tasks the network had to respond with the second cue or the opposite of the second cue, respectively. All context-cue pair contingencies are detailed in **Fig. 1b**.

Preceding Cue A was a foreperiod of 200 ms in which only the rule input was on. The rule input is presented throughout the trial. Following the foreperiod, one of two Cue A channels was set to 1 for 100 ms. Next both Cue A channels were returned to 0 for a delay period of 2000 ms. After the delay one of the two Cue B channels was set to 1 for 100 ms. Following Cue B onset the network was instructed to produce a steady output of 1 in the output channel corresponding to the correct response for that trial. Output was masked until 100 ms following Cue B, such that output magnitudes prior to that time were not considered in the loss function (and therefore unconstrained).

### RNN training

The RNN’s hidden (recurrent) weights were initialized as a random gaussian matrix with spectral radius equal to 1.1, and the initial input and output weights were initialized with the Xavier method^52^. RNNs were trained using an Adam optimizer^53^ as implemented in TensorFlow 1.11.0, and with regularization on weights and unit activity^16,54^. Input, recurrent, and output weight matrices were regularized with L1 penalties of 0.01, and unit activity was regularized with an L2 penalty of 1. The overall loss function is thereby given by:

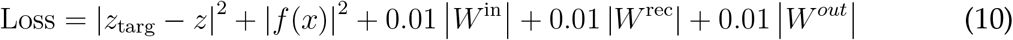

where *z*_targ_ is the target output.

The task was trained and tested with noise injected into input channels and recurrent units. Gaussian noise (*μ* = 0, *σ* = 0.01) inserted into each stimulus channel, and into each recurrent unit (*μ* = 0, *σ* = 0.1), at each time point.

Networks were trained with a curriculum learning regime in which the memory delay duration was progressively extended over training^48^. Initially the network was trained for 2,000,000 trials with a delay duration of 400 ms. This was followed by training the network for 200,000 trials on increasing delays of 200 ms greater than the previous iteration. When the delay length reached 2000 ms the network was trained again for a final 2,000,000 trials. Training proceeded with batches of 128 trials.

To test for robustness of results in RNNs across initial conditions and training histories, 20 replicates of the network were trained, each starting from a different random initialization.

### RNN Analysis

#### Explained variance analysis

We utilized a linear model with binary coded contingency as predictors to identify the proportion of across-trial state variance explained by each contingency for each unit *j* in our network:

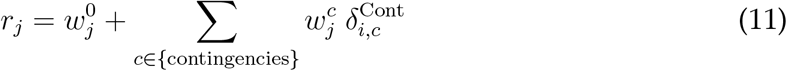

where, as in the empirical analysis, 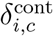 is 1 when the contingency of trial *i* matches contingency *c* and 0 otherwise. This regression model thereby uses 5 parameters per unit (1 baseline and 4 contingency). The proportion of explained variance can be calculated per unit *j*: EV_*j*_ = 1 − SSE_*j*_/SST_*j*_.

We defined the late delay epoch state as the dynamic state variable of each unit at the time step prior to onset of Cue B. For demonstration of an example unit, **Fig. 5a** plots the mean rectified state variable, averaged across trials for each rule and Cue A condition, for the recurrent unit that had the most [0-1] contingency variance.

In order to measure differences in sample and delay tuning we utilized linear models to calculate, separately for each unit, the amount of state variance across trials explained by Cue A identity and contingency. For **Fig. 5c**, variance explained was calculated at two time points: the final time point before Cue A offset (Sample) and the time point prior to Cue B onset (Delay). Explained variance was calculated as below, and plotted is the average variance explained across units for each feature and time point.

Model 1 used contingencies (Eq. 11). Model 2 used Cue A identities:

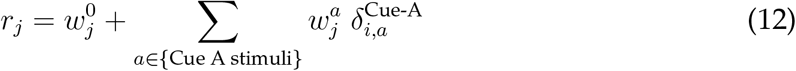

The explained variance for given feature *f* (contingency or Cue A) is given by:

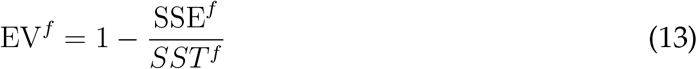

These regression models and calculation of explained variance can be performed in a time-resolved manner to characterize the time course of coding in the network (**Fig. 5b,c**).

For comparison to the experimental results from Rainer et al. (1999)^6^, we utilized the WebPlotDigitizer^55^ to replicate plotted data values (**Fig. 5d**).

#### Dynamic coding axis

For **Fig. 5e**, at each time point, and separately for each rule, we calculated the first principal component (PC) of network states across trials, following prior experimental studies^22,39,40^. We then measured the angle between first PCs from each pair of time points identifying the degree to which the dominant coding axis had shifted between time points^22^. We then plotted the angular similarity between time points averaged across rules.

#### Dynamic decoding

For **Fig. 5f**, we first separated the data into two equal halves of 500 trials as train and test sets. We trained a linear kernel support vector machine (SVM), for each rule to classify Cue A on the stimulus epoch activity of the train set (sample decoder in **Fig. 5f**), and then tested it on each time point throughout the task on the test set. Next we trained a separate linear SVM to classify Cue A at each time point on the train set, testing it on the test set trials from the corresponding time point (dynamic decoder in **Fig. 5f**). This analysis is analogous to dynamic decoding analyses in prior experimental studies examining cross-temporal generalization of working memory representations in prefrontal cortex^23,24,38^. Plotted is mean SVM accuracy averaged across rules.

#### Contingency subspace

We determined the first axis of the contingency subspace via a linear regression to identify an axis which could decode the expected response to an upcoming Cue B=0, based on the Cue A and rule on that trial, using the late delay epoch unit activation. We similarly identified the second axis through a regression for the expected response to an upcoming Cue B=1. While axes were determined independently, we found they were always nearly orthogonal over 20 replicate RNNs with a mean between axis angle of 92.6 degrees (sd 4.7 degrees). Further the regression method caused a modest rescaling of the original space with a mean axis norm of 1.4 (sd 0.17).

To get the contingency representation we projected the state vector from individual trials into the plane identified. To avoid any issues with overfitting, we fit the regression on half our data which we used as a training set, and utilized the held-out half of our data for figures and analysis.

#### Variance captured by subspaces

In order to test the hypothesis that more condition-wise variance is explained by the contingency subspace than would be expected by chance, we examined whether random two dimensional subspaces would contain equal variance (**Fig.= 4c**). First we generated random orthogonal planes using Gram-Schmidt orthogonalization and rescaling, normalized such that the axes were equal in norm to the contingency subspace axes. We then orthogonalized the two contingency subspace axes and measured the sum of the state variance in the contingency plane (the red vertical line). We compared the variance in the contingency subspace to the state variance across the 20,000 randomly sampled orthogonal planes (**Fig. 4c**). We then ran this analysis over our 20 replicate networks and calculated a normalized variance measure, defined as the variance in the contingency subspace divided by the mean of the variance across random orthogonal planes (**Fig. 4d**).

#### Perturbation analysis

To functionally identify the relevance of the contingency subspace for behavior we applied a state-space perturbation analysis. A random vector was added to the state of the network at the time point immediately preceding Cue B onset. This perturbation was either linearly dependent on the axes of the contingency subspace and therefore lie within the subspace, or orthogonal to the subspace. We tested random perturbations of increasing vector norm (magnitude) and measured the resulting decrease in accuracy. We repeated this analysis for our 20 replicate RNNs. Plotted are lines for the mean accuracy across replicates for a given perturbation either in the subspace or out of the subspace with the band representing the 90% confidence interval (**Fig. 4e**, **Supplementary Fig. 4**).

### Singular value decomposition (SVD) analysis of CDL task

First we divide the CDL task into (i) pre-delay features, i.e., features known to the agent prior to the delay, and (ii) post-delay features. Pre-delay features include the rule and Cue A, while post-delay features include the Cue B and response. We represent each feature as a one-hot encoding. Then we can build an interaction matrix for Cue A × Rule and a separate interaction matrix for Cue B × response. Finally we construct the correlation matrix between our two interaction matrices, and factorize it with a singular value decomposition (SVD). This results in a *U* matrix which represents loadings for our task conditions, a *V* matrix that represents loadings onto our contingencies, and a Σ matrix that represents the weighting of those loadings.

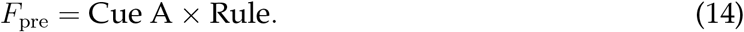

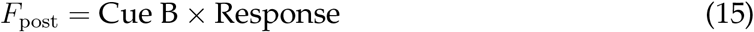

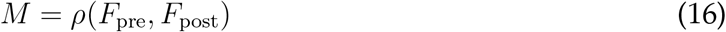

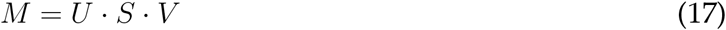

Results of this SVD analysis are shown in **Supplementary Fig. 1**.

### Task pair analysis

To identify Cue A tuning in the contingency subspace we first mapped each Cue A × Rule condition of a given pair of tasks to its associated contingency state. Here we use an idealized set of orthonormal contingency coding axes which provide the only tuning in the system, with the first axis encoding the response if Cue B=0, and a second axis encoding the response if Cue B=1 (**Fig. 6a**).

Each task pair is represented four possible states, 2 Cue A states × 2 rule states. For each pair we first averaged states across rules. We took the measured Euclidean distance between Cue A=0 mean and the Cue A=1 mean as our theoretical prediction of cue tuning. Similarly, to identify rule tuning we averaged across Cue A states. The Euclidean distance between the means for the two rules in the contingency subspace was taken as our theoretical prediction of rule tuning. We repeated this procedure through all 100 possible pairs of rules (including each rule paired with itself) to generate the full matrix of predictions. We then defined a theoretical cue:rule tuning ratio as the ratio of our predicted cue tuning over predicted rule tuning (**Fig. 6b**).

In each of 20 replicate RNNs trained on the CDL task, we isolated trials belonging to a pair of tasks. We then averaged the state vectors by condition (Cue A × Rule pairs). As above we then averaged the state of the network and measured the Euclidean distance in state space across rule to get cue tuning and across cue to get rule tuning. We repeated this procedure for all task pairs as above. We then divided our measured cue tuning by rule tuning to get a RNN rule to cue tuning ratio. Finally we averaged across our 20 replicates. To compare the RNNs to theoretical predictions, we used a Spearman rank correlation test to identify similarities in the pattern of the lower triangular matrices of both the predicted and measured cue to rule tuning ratios (**Fig. 6c**).

### Comparison models

#### RCN model

We implemented a randomly connected network (RCN) model^8,25^ to investigate differential predictions between unstructured-high dimensional nonlinear representations and our task-optimized RNN models. To generate internal representations for RCN models, we generated one-hot representations of rule, Cue A, and Cue B for each trial, which we term *u*_inst_ of dimensionality 1000 × 14. Then we multiply these representations by a matrix, *W*_rand_, of dimension 14 × 200, the number of input variables by the maximal dimensionality of the 200 recurrent units of the RNN. Elements of *W*_rand_ were drawn from a random normal distribution. A threshold variable *θ* for each element drawn uniformly from the interval [−1, 1] were then subtracted. Finally we transform the representation with a sign operation such that values are represented by ±1.

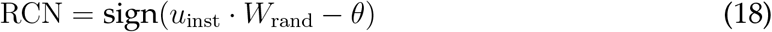

For the inputs model we took the one-hot representations of rule, Cue A and Cue B, *u*_inst_, and simply ran analysis directly on these representations.

#### Representational similarity analysis (RSA)

To compare our trained RNN to the RCN model in representational structure, we applied a representational similarity analysis to both networks^27^. For the RNN we took the activity vector at the time point prior to Cue B onset. We generated representational similarity matrices by averaging activity across trials for each Cue A × rule condition, and then taking the pairwise Pearson correlation between the averaged activity vectors (**Fig. 7a,b**).

We then compared these matrices to three binarized candidate representational matrices: (i) Cue, with two conditions being similar if they had the same Cue A; (ii) rule, with two conditions being similar if they shared a rule; and (iii) contingency, with two conditions being similar if they share a contingency (**Fig. 7c**). We repeated this analysis for 20 randomly constructed RCNs and our 20 replicate RNNs. Plotted is the distributions over these replicates separated for cue, rule and contingency, respectively (**Fig. 7d**).

#### Variance explained

In order to identify the proportion of variance explained by Cue A, rule, and contingency in each model, we fit a linear model for each feature to each unit states across trials using a regularized least squares (*λ* = 0.1). For the RNN, we used the state at the time step prior to Cue B onset. We then calculated the proportion of variance explained as 1 − SSE/SST. We then conducted this analysis for each unit, taking the average over a given network. Shown is the distribution of mean unit variance explained by that feature over 20 RCN replicates and 20 RNN replicates (**Fig. 7e**).

#### Dimensionality

To measure effective dimensionality of states in our models, we used the participation ratio^30,56^ calculated as 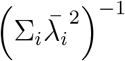, where 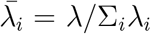 and {*λ_i_*} are the eigenvalues of the covariance matrix. We applied this measure to the RNN states at time step prior to Cue B onset. We also used this analysis to measure the effective dimensionality of the RCN activity and of the input data itself as comparisons (**Fig. 7f**).

#### Cross-context generalization

We utilized a cross-condition generalization (CCG) analysis^32^ to test the extent to which the geometry of the contingency subspace represented common information between different trial conditions with the same contingency state. We fit the contingency subspace, as described above, except we held out all trials with a given rule–Cue A condition. We then projected the held-out trials into the subspace and used the four quadrants of the contingency subspace as a classifier, with each trial being classified as the contingency state of the appropriate quadrant. We repeated this analysis for our 20 RNN replicates, as well as for our 20 RCN replicates. Plotted is the distribution of accuracies for the CCG classifier across networks (**Fig. 7g**).

### Partitioned variance analysis

To measure the extent to which contingency was over-represented in our RNN model compared to statistically matched nonlinear structure, we implemented a permutation test and identified mean unit state variance explained. Then we permuted contingencies between Cue A × rule pairs to generate pseudo-contingencies (e.g. all Cue A=0, Rule=AND trials were assigned the same new random contingency) (**Fig. 8a**). We regressed out linear effects of Cue A and rule, fitting a linear model and taking the residual for further analysis. Then we used our pseudo-contingency labels to fit a linear model using regularized least squares (*λ* = 0.1). Finally we calculated variance explained as 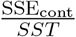. We compared the actual contingency variance explained to 1000 randomly sampled pseudo-contingency sets (**Fig. 8b**).

Specifically, we perform the following regression model:

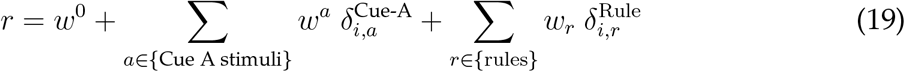

We define the residuals of this regression model:

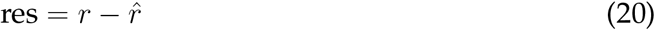

where 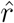 is the predicted activity for each trial. We next perform a second regression on the residual:

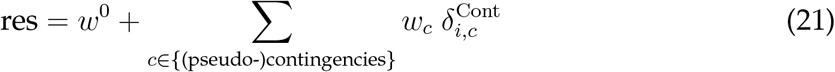

using either the true contingencies, or the shuffled pseudo-contingencies to generate the null distribution.

We repeated this analysis for the RCN data, as well as all replicates of the RNN (**Fig. 8b,c**). We calculated normalized variance as the mean variance explained by contingency labels over the mean variance explained by pseudo-contingency permutations. Shown is the distribution across RNN replicates (**Fig. 8d**).

### Code and data availability

Task code, behavioral data, trained models and analysis will be made freely available in public repositories upon publication.

---

**Supplementary Figure 1:**
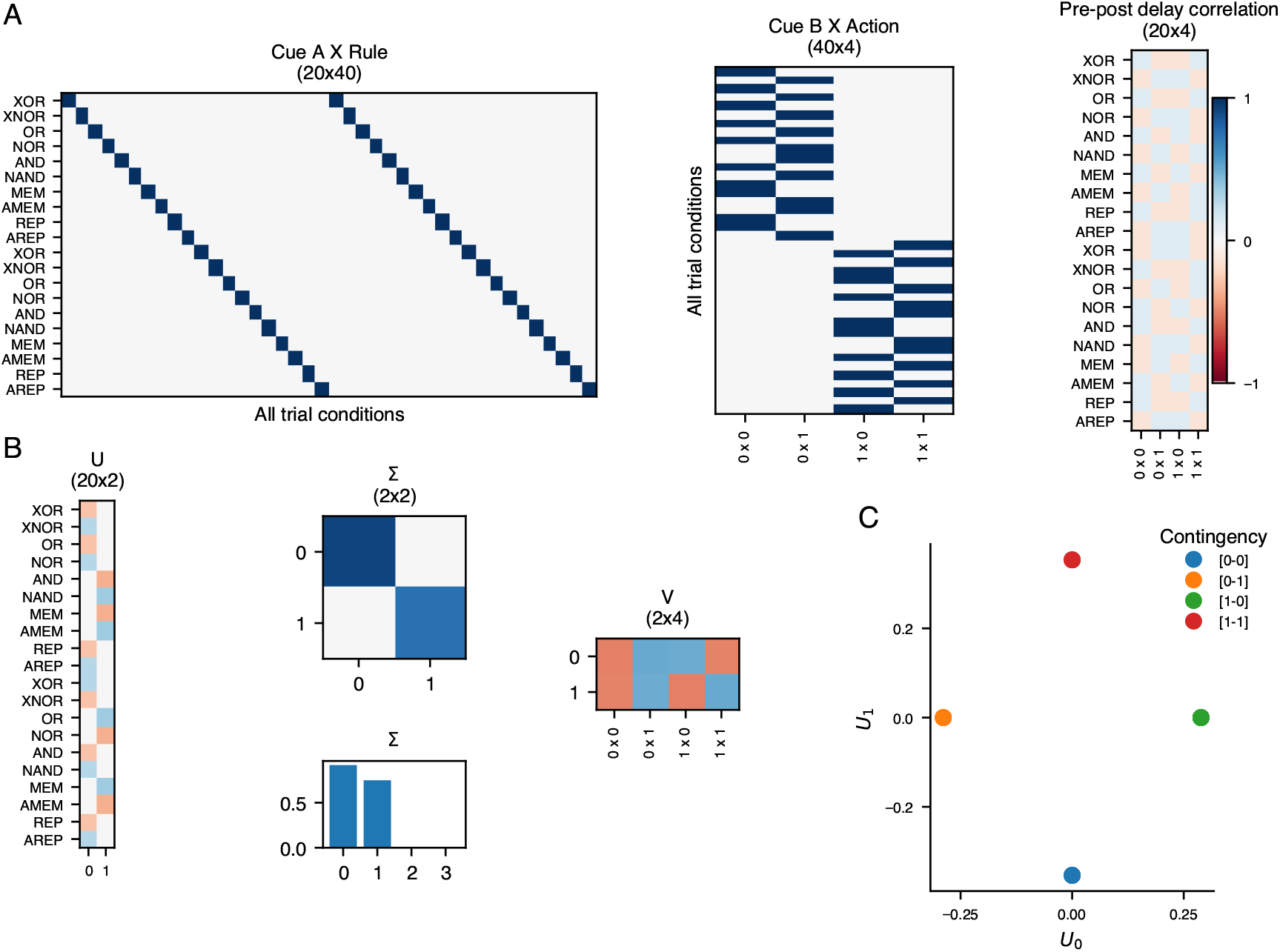
Singular value decomposition (SVD) analysis of CDL task. **(a)** Left: Cue A by Rule interaction matrix (20 × 40), representing the interaction of all trial information provided prior to the delay, Middle: Cue B by Action interaction matrix (40 × 4), the trial information delivered post delay, Right: the Pearson correlation between pre-delay and post-delay features. **(b)** SVD of the correlation matrix in A, into a diagonal matrix Σ, and two orthogonal matrices U and V, representing a basis in pre-delay and post-delay features respectively. The bar graph represents the fraction of variance explained by each mode of the decomposition, **(c)** A projection of U values colored by the contingency of the associated trial condition. SVD of the CDL problem naturally divides along contingency into a two-dimensional subspace.

**Supplementary Figure 2:**
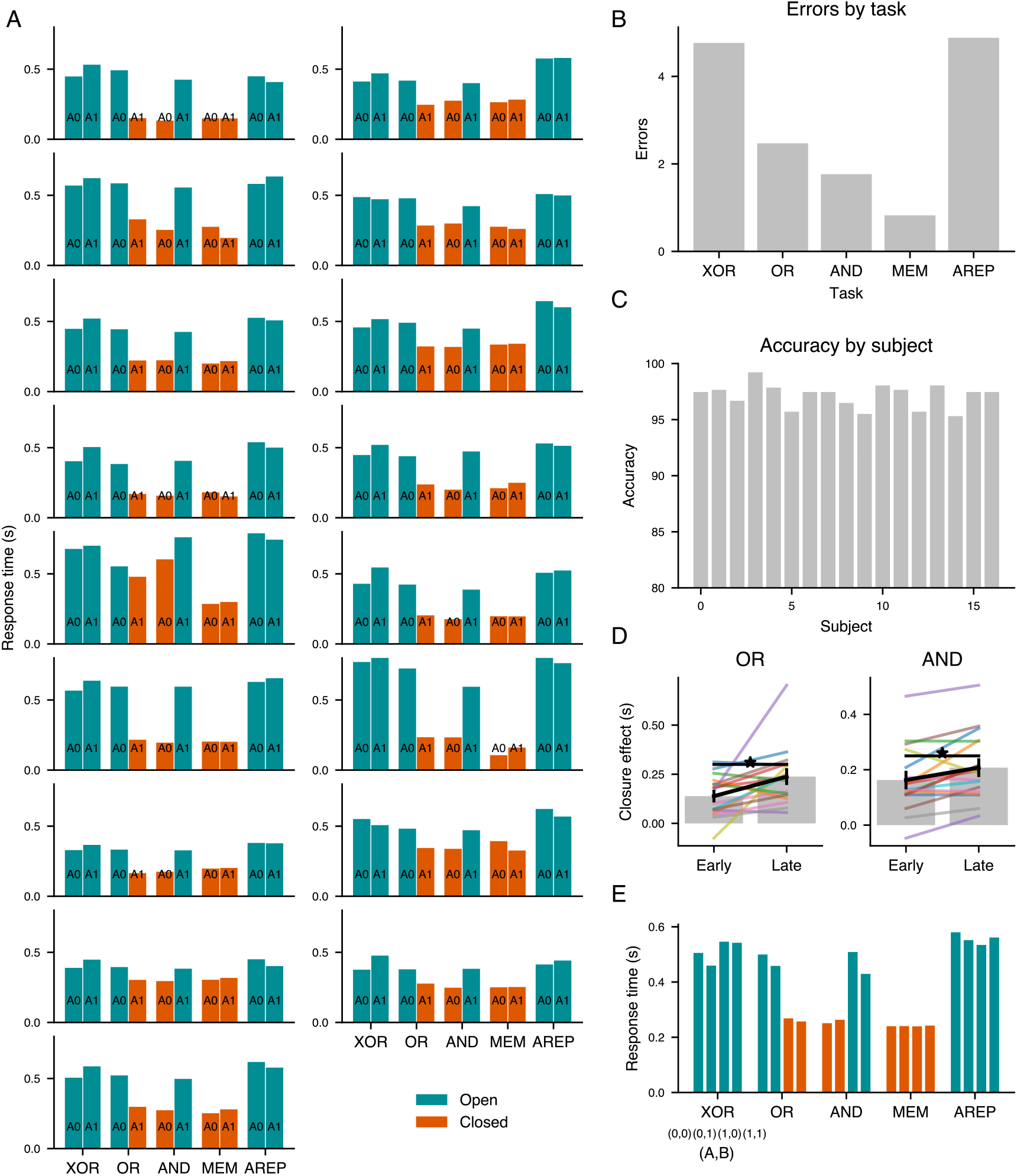
Individual participant behavior on CDL task. **(a)** All participant data plotted as in **Fig. 3b** with mean reaction time averaged across Cue A and rule pairs. **(b)** Mean errors per task averaged across participants. **(C)** Overall accuracy presented for each participant **(D)** Individual participant data for change in closure effect. Each line represents a single participant, with bars covering the mean early (first two blocks) and late (last two blocks) session closure effect. Black line represents the average change from early to late. **(E)** Mean accuracy by condition, averaged across participants, divided by rule, Cue A and Cue B combination.

**Supplementary Figure 3:**
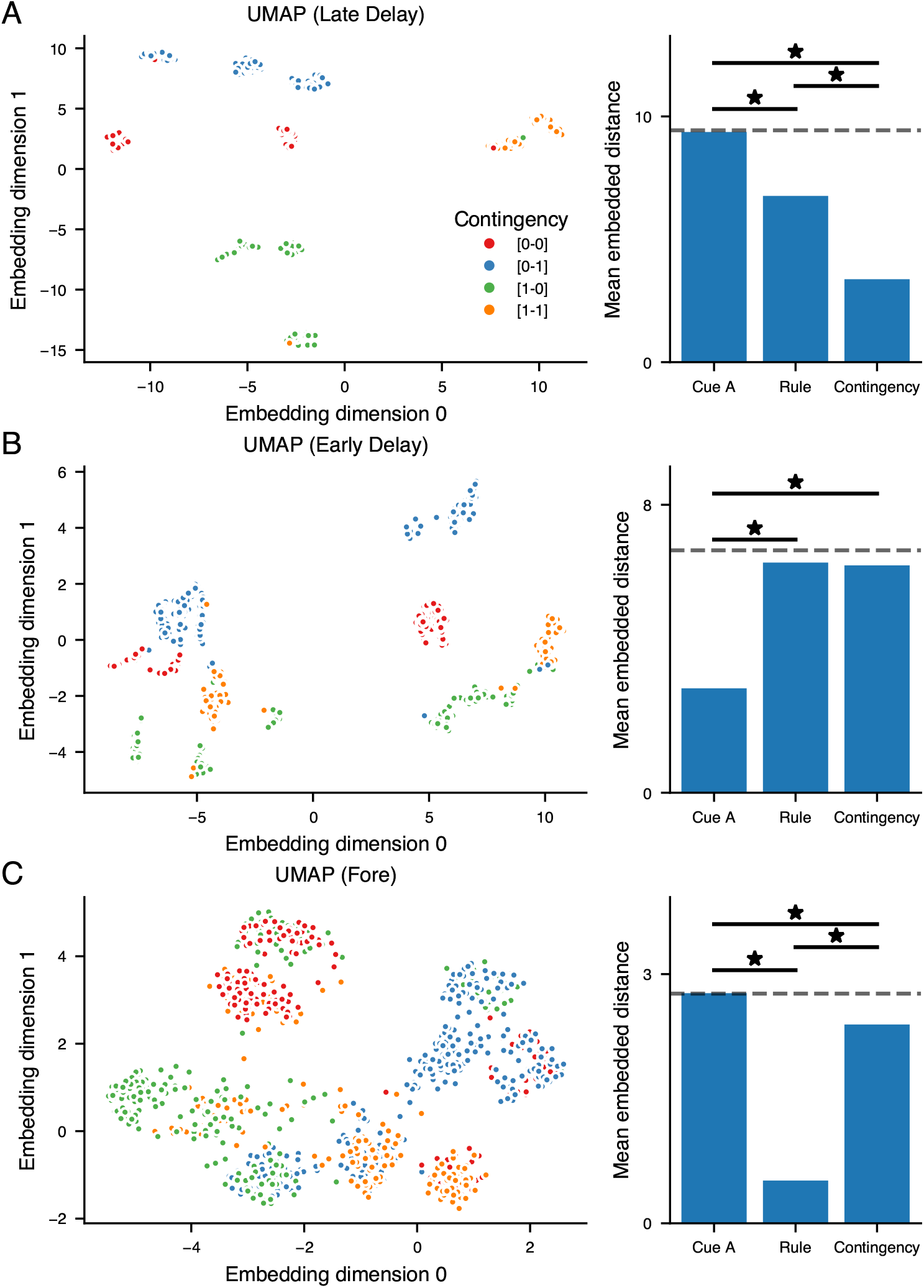
UMAP analysis of RNN representations. **(a)** Left: UMAP (Uniform Manifold Approximation and Projection) performed on states of the network in the late delay epoch (time point prior to Cue B onset) colored by contingency. Right: Quantification of embedding distance. For each feature first we averaged the embedding values for all trials by condition across that feature (e.g. Cue A=0 and Cue A=1 for Cue A). Then we calculated the mean Euclidean distance from each trial embedding to the its associated centroid. UMAP embeddings in the late delay epoch clustered more substantially by contingency than by Cue A (one-way anova, p<0.001) or rule (p<0.001). Dashed line represents chance embedding distance, generated through an analysis of distance to centroids of randomly partitioned trials. **(b)** Analysis as in **(a)** with activity states from the early delay epoch (200 ms after Cue A offset). We found that in the early delay epoch, Cue A centroids significantly outperformed rule (p<0.001) and contingency (p<0.001) organization. **(c)** Analysis as in **(a)** with activity states from the fore epoch (50 ms before Cue A onset). In the foreperiod UMAP embeddings were more highly organized by rule than Cue A (p<0.001) and contingency (p<0.005). Despite this there remains significant contingency organization during the foreperiod, with lower contingency embedding distance than Cue A (p<0.001).

**Supplementary Figure 4:**
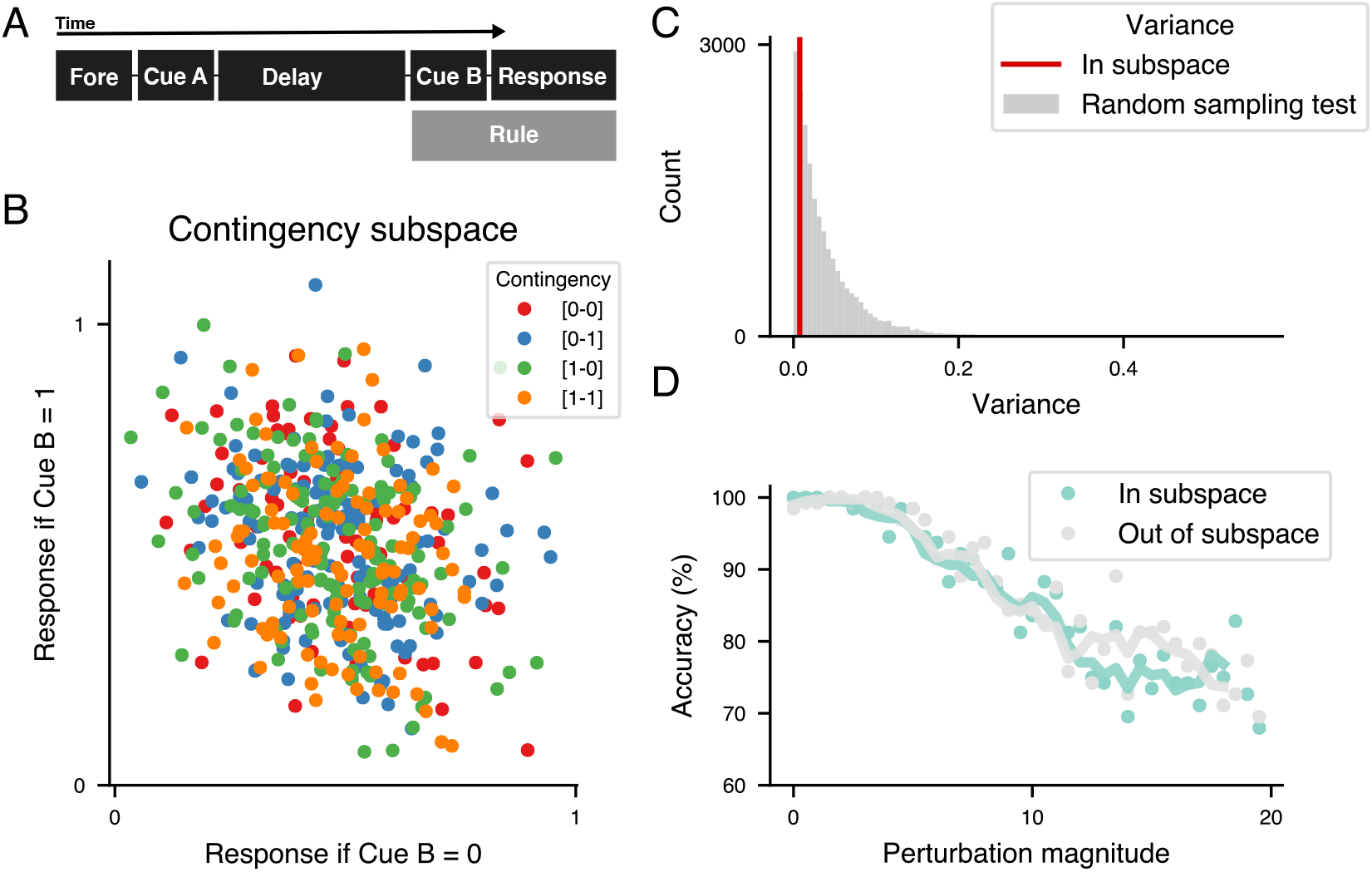
Late rule task. **(a)** A schematic of the late rule task. All events in the trial proceed as described for the original task, with the exception that rule input only onsets at the start of Cue B onset rather than being tonically on throughout the trial. This makes calculating contingency during the delay impossible. **b)** The contingency subspace as calculated in **Fig. 4b** for the late delay network shows no substantial organization by contingency. **(c)** The contingency subspace in **(b)** shows no preferential variance as compared to randomly chosen subspaces. **(d)** Despite showing no contingency subspace, performance of the network is high with near 100% accuracy in the unperturbed model. Further validating that a contingency solution is not being utilized by this network, the late rule network shows no preferential vulnerability to perturbations in the contingency subspace as opposed to orthogonal to it, for any perturbation magnitude.

**Supplementary Figure 5:**
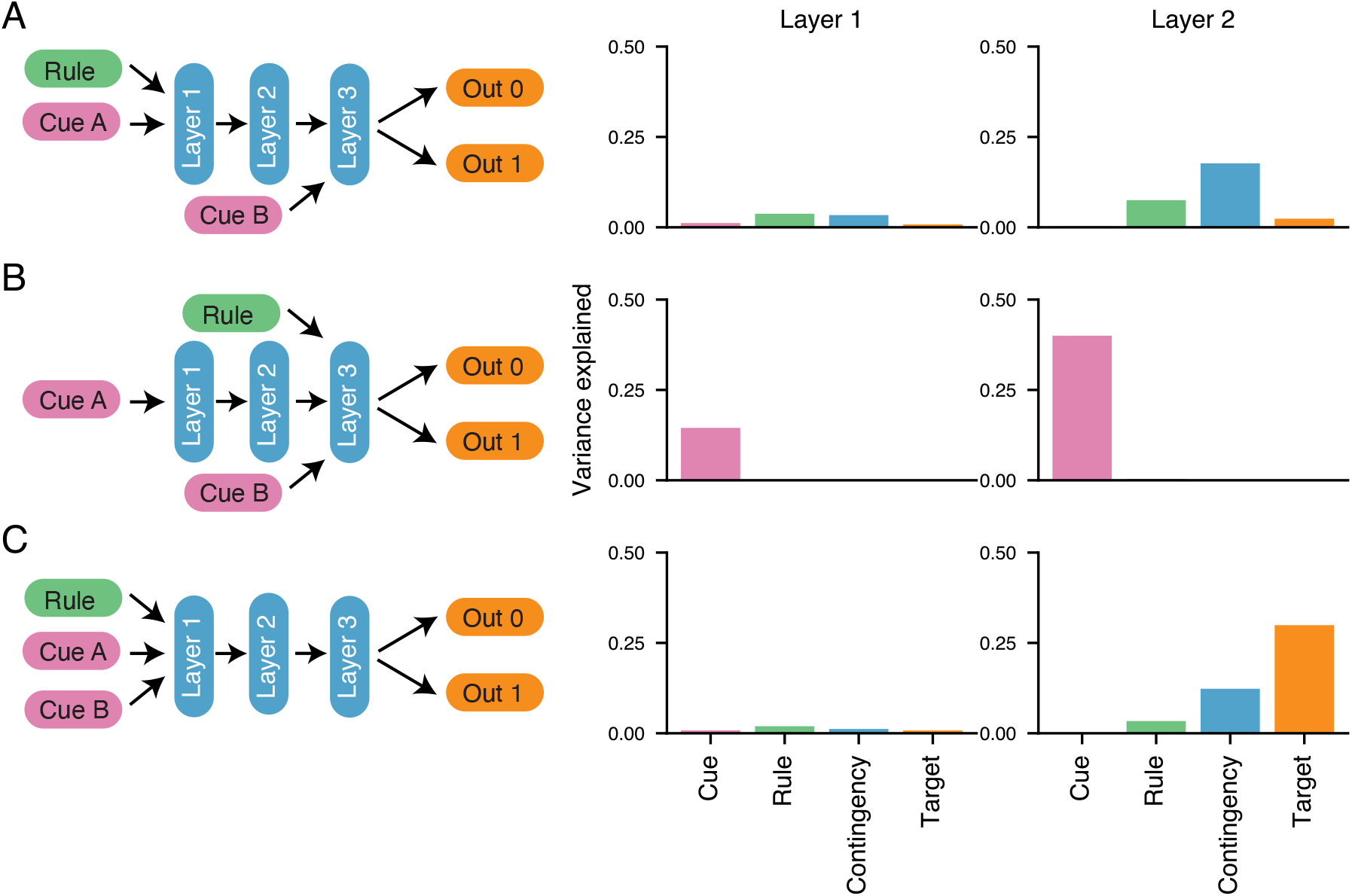
Feedforward neural networks and stepwise computation. A feedforward model of the CDL task using a 3 layer fully connected neural network with ReLU nonlinearities. Left: Schematic of input regime. Inputs were represented as in the RNN by one hot vectors, with dimensionality 2 for Cue A and Cue B and dimensionality 10 for the rule input. Each hidden layer was composed of 200 ReLUs. All weights were initialized with Glorot initialization. Networks were trained for 20,000 iterations using the Adam optimizer (learning rate=0.001) on a mean squared error loss function with L2 regularization (*λ*=1) on hidden unit activity. Right: The mean unit state variance explained across trials by each feature (Cue A, rule, contingency and target) for layers 1 and layers 2 of the model. **(a)** As in the CDL task, Cue A and rule are input into the first layer and Cue B is input into the third layer. Information processing of the first layer roughly approximates the early delay, while the second layer represents the late delay and third layer the post delay epochs. Units in the layer 2 (late delay) but not layer 1 (early delay) are best explained by contingency as in the recurrent network. **(b)** As in the late rule network, only Cue A is input into the first layer, with both rule and Cue B being input into layer 3. The network as in the recurrent network demonstrates only Cue A variance in both layers 1 and 2. **(c)** All input, rule, Cue A and Cue B, enters at the first layer. In no layer does contingency best explain unit variance.

**Supplementary Figure 6:**
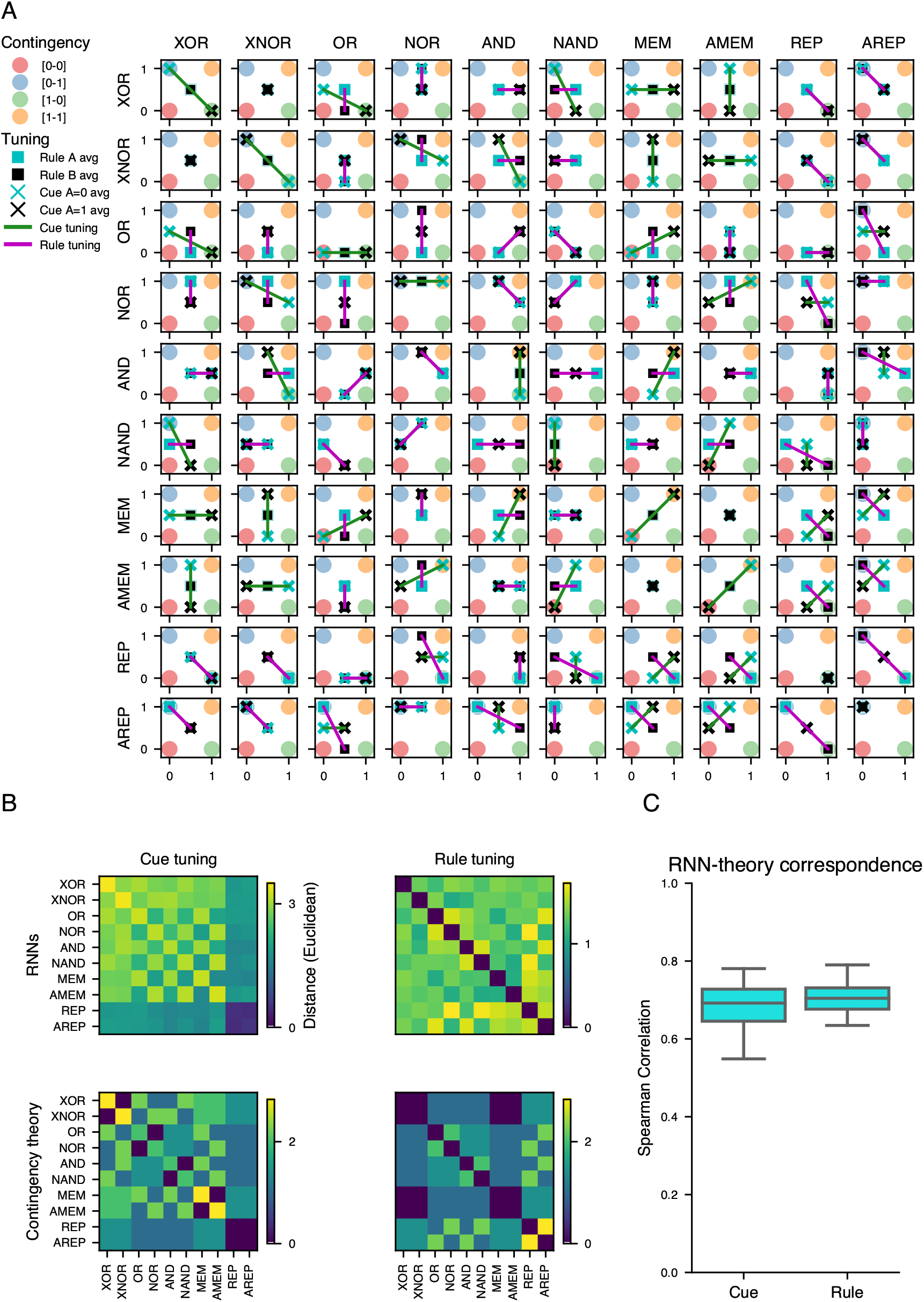
Task pair tuning analysis. **(a)** Representation for all 55 task pairs of predicted theoretical contingency induced tuning for rule and Cue A. Circles, colored by contingency, represent the centroid for trials of each contingency in the contingency subspace. Cyan(Black) Xs represent the location of trials averaged across rules for Cue A=0(Cue A=1). Cyan(Black) Xs represent the location of trials averaged across rules for Cue A=0(Cue A=1). Cyan(Black) squares represent the location of trials averaged across cues for Rule A(Rule B), with Rule A on the x-axis and Rule B on the y-axis. Green lines represent the direction and magnitude of cue tuning induced by representation in the contingency subspace. Magenta lines represent the magnitude and direction of rule tuning induced by representation in the contingency subspace. **(b)** Measured and theoretical tuning for Cue A and rule. Measured tuning represents the Euclidean distance between the trial averaged recurrent unit states from the late delay epoch. Theoretical tuning is the Euclidean distance between representations averaged, within Cue A or rule for cue tuning and rule tuning respectively, in the contingency subspace. **(c)** Spearman correlation calculated across 20 replicate RNNs between theoretical and measured Cue A and rule tuning. Correlation was measured between lower triangular elements between RNN tuning matrices and contingency theory matrices separately for cue and rule tuning.

